# SoMAS: Finding somatic mutations associated with alternative splicing in human cancers

**DOI:** 10.1101/2023.07.06.547933

**Authors:** Hua Tan, Valer Gotea, Nancy E. Seidel, David O. Holland, Kevin Fedkenheuer, Sushil K. Jaiswal, Sara Bang-Christensen, Laura Elnitski

**Affiliations:** Genomic Functional Analysis Section, TFGB, NHGRI, NIH, Bethesda, MD

## Abstract

Aberrant alternative splicing is prevalent in cancer and affects most cancer hallmarks involving proliferation, angiogenesis, and invasion. Somatic point mutations can exert their cancer-driving functions via splicing disruption. We propose “SoMAS” (**So**matic **M**utation associated with **A**lternative **S**plicing), an efficient computational pipeline based on principal component analysis techniques, to explore the role of somatic mutations in shaping the landscape of alternative splicing via both *cis*- and *trans*-acting mechanisms. Applying SoMAS to 33 cancer types consisting of 9,738 tumor samples in The Cancer Genome Atlas, we identified 908 somatically mutated genes significantly associated with altered isoform expression in three or more cancer types. These genes include many well-known oncogenes/suppressor genes, RNA binding protein and splicing factor genes with both biological and clinical significance. Many of our identified SoMAS genes were corroborated to affect gene splicing by independent cohorts and/or methodologies. With SoMAS, we for the first time demonstrate the potential network of somatic mutations associated with the overall splicing profiles of cancer transcriptomes, bridging the genetic and epigenetic regulation of human tumorigenesis in an innovative way.

## Introduction

Alternative splicing (AS) is a widespread biological process responsible for most of the transcript structural variation and proteome complexity and diversity in human genomes [1]. Aberrant AS is common in human cancers and affects cancer progression in many aspects known as the ‘hallmarks of cancer’ [2–4]. Somatic disease-causing mutations, or driver mutations, are commonly assumed to exert their effects by altering amino acid sequences, which eventually impairs the normal function of a protein [5–7]. However, increasing evidence confirms that somatic mutations in both coding and noncoding regions [8] can dramatically disrupt gene splicing (referred to as splicing mutations) through a variety of effects including exon skipping, intron retention and frameshifts [9–12].

Several studies have identified the importance of somatic mutations, primarily point mutations, or single nucleotide variants (SNVs), on AS in human cancers. For example, a study of 1,812 patients from six cancer types using TCGA (The Cancer Genome Atlas) database identified ∼900 somatic splicing mutations in coding regions, of which 163 SNVs likely caused intron retention or exon skipping [10]. This work further demonstrated an enrichment of SNVs causing intron retention in tumor suppressor genes (TSGs), and highlighted intron retention as a common mechanism of TSG inactivation. A more comprehensive study known as MiSplice (mutation-induced splicing) utilizing ≥8,000 tumor samples across all 33 TCGA cancer types characterized 1,964 somatic mutations with evidence of creating alternative splice junctions within windows of ±20 bp from each mutation (i.e., *cis*-acting), termed “splice-site-creating mutations” (SCMs) [13]. Neoantigens induced by SCMs proved to be more immunogenic than missense mutations, making them good candidates for immune therapy. Another pan-cancer association analysis of genetic variation and AS termed sQTL (splicing quantitative trait loci) based on genomics data from the same 33 TCGA cancer types identified 32 *cis*- and seven *trans*-acting sQTLs (termed *cis*-sQTLs and *trans*-sQTLs, respectively), which included those on the well-known splicing factors SF3B1 and U2AF1 [14].

In addition to the abovementioned general and integrative mutation-splicing association studies, other research addressing somatic mutations in known splicing factors (SFs) and regulators has confirmed the *trans*-acting effects of SNVs on gene splicing, and their clinical relevance through either computational or experimental approaches [15–17]. Together, these studies well established that somatic SNVs that disrupt splicing represent a major pathogenic mechanism in human cancers.

Despite the many merits of the existing studies, methodological limitations can easily miss sQTLs and SNV-associated AS events. In particular, the existing pipelines and models are invariably limited to a direct, one-to-one mapping between an SNV and a specific, mostly local AS event. Specifically, in search of *cis*-sQTL, existing studies restricted their search to the flanking exons or neighboring exon-intron junctions; whereas the search for *trans*-sQTLs considered only known SF and splicing regulator genes. These strategies focused on outcomes of direct regulation of splicing by mutated SFs or regulators but ignored SNVs occurring in many other RNA-binding proteins (RBPs) and co-regulators, especially those regulators upstream of splicing/transcription machinery (e.g., transcription factors or TFs) that work in an indirect way. A striking example of indirect regulation is seen in the effects of mutant TP53 on RNA splicing via upregulating an intermediate RBP called hnRNPK [18]. The direct gene-by-gene association study has two methodological drawbacks: (i) it necessitates a huge number of SNV-AS pairs in order to cover all possible combinations, incurring an intense computational burden, especially for a pan-cancer analysis; and (ii) the statistical power of this direct gene-by-gene correlation is diminished due to the generally low mutation frequency of most cancer driver genes [19, 20], which requires a sophisticated multiple testing correction to mitigate the bias and to accommodate different cancer types.

To address these challenges and methodological gaps, we developed a computational pipeline called SoMAS (Somatic Mutation associated with Alternative Splicing) that integrates matched DNA-seq and RNA-seq data of a particular cancer type to explore the impact of somatic SNVs on RNA splicing profiles. We showed that both RNA-seq (for gene isoform expression) and DNA-seq (for gene somatic mutation) data in TCGA support a novel molecular taxonomy of tumors in addition to the current tissue histology-based pathology, and hence provide us a great opportunity to investigate the association between gene somatic mutation and gene isoform expression patterns. Instead of examining the direct, one-to-one association between each SNV and a specific AS event, SoMAS measures the association between each eligible SNV (with mutation frequency exceeding a designated threshold) and the overall expression pattern of the whole transcriptome consisting of ∼73K annotated transcripts, using the principal component analysis (PCA) technique. Eventually, with SoMAS we detected a total of 908 SoMAS genes that potentially impact gene splicing (quantified as gene expression at the isoform level) in at least three cancer types. These pan-cancer SoMAS genes include many well-recognized oncogenes, tumor suppressors and RBPs/SFs/TFs, many of which were verified to affect splicing of predicted targets by previous studies with independent cohorts and/or methods. We further characterized the *cis*- and *trans*-effects of those SoMAS genes and confirmed their biological and clinical significance with a series of functional analyses. SoMAS is the first tool that systematically explores the implications of somatic SNVs in genome-wide splicing and transcription regulation across pan-cancer and proves to be efficient and versatile.

## Results

### Gene isoform expression and somatic mutation profiles provide new molecular taxonomy beyond histopathological classification

To study the association between gene mutation and gene isoform expression, we examined matched RNA-seq and DNA-seq (whole-exome) data from The Cancer Genome Atlas (TCGA) (Figure 1A). The RNA-seq data measured the RNA abundance of 73,599 isoforms derived from 29,181 genes (including 20,429 coding and 8,752 noncoding genes) (Table S1). The DNA-seq data detected somatic mutations on 22,029 genes. These data were collected from a total of 10,464 cancer samples (including 9,738 tumor samples and 726 tumor-adjacent normal tissue samples) across 33 cancer types (ranging from 45 to 1,215 samples per cancer type).

**Figure 1.**
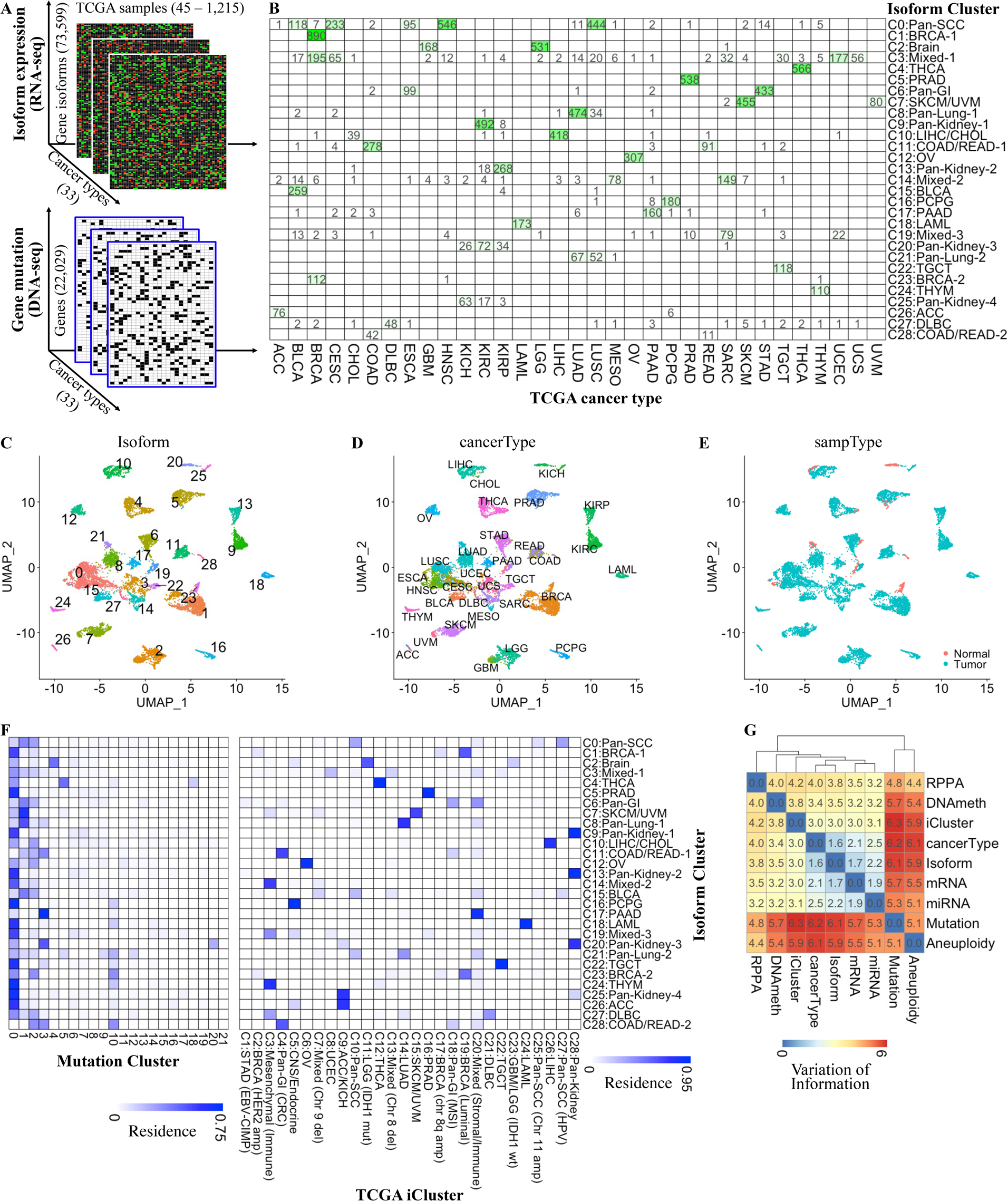
Gene isoform expression and somatic mutation profiles provide new molecular taxonomy beyond tissue-histology classifications. (A) TCGA multi-omics data employed in the study include matched RNA-seq for isoform expression (upper) and whole-exome-wide DNA-seq data for gene somatic mutation (lower) across 33 cancer types. (B) Cluster residence heatmap shows the number of samples from a given cancer type that reside within each of the 29 (0 – 28) annotated Isoform clusters. (C) The UMAP visualization of 10,464 TCGA cancer samples based on expression of 73,599 isoforms. Each dot represents a TCGA sample and is colored/marked by the 29 Isoform clusters. (D-E) Samples on the same Isoform UMAP map are colored by TCGA cancer types (D) and tumor status (E). (F) Cluster residence heatmap shows the percent of each Isoform cluster that overlaps with each Mutation cluster (left) and TCGA iCluster (right). (G) Variation of information analysis of clustering schemes derived from various TCGA data types. The cluster membership of Aneuploidy, mRNA, miRNA, RPPA, DNA methylation (DNAmeth) and iCluster were derived from (Hoadley et al., 2018). The membership of Isoform was determined in this study. See also Table S1 and Figure S1.

Clustering analysis based on gene isoform expression (normalized RSEM values) across all 33 cancer types (see Methods) identified 29 distinct “Isoform” clusters (labeled as C0–C28, Figure 1B). We annotated these clusters according to the observed cancer type enrichment. For instance, cluster C2 was annotated as Brain because most samples in this cluster were from brain tissue-originated cancers: GBM (Glioblastoma) and LGG (Lower-grade Glioma). The 29 molecular subtypes generally well reflected the tissue origin of samples in each cluster (Figure 1B-D), and discernibly distinguished the tumor and normal samples (Figure 1E). In addition, we observed several intriguing patterns from the clustering results. For example, samples from the same or adjacent organ/tissue tended to cluster together, such as the Pan-GI (gastrointestinal) cluster (C6), the COAD/READ (large intestine) clusters (C11 and C28), the Pan-Lung clusters (C8 and C21), and the Pan-Kidney clusters (C9, C13, C20 and C25). Additionally, tumors originating from the same cell type were likely to show similar isoform expression pattern, such as the Pan-SCC (squamous cells) cluster (C0) and SKCM/UVM (pigment-producing cells) cluster (C7). Some cancer types split into two or more distinct clusters such as the breast (BRCA) and esophageal (ESCA) cancers. We also detected three mixed clusters with samples from multiple cancer types (C3, C14 and C19).

Clustering analysis of the same TCGA tumor samples based on whole-exome somatic mutation data (see Methods) detected 22 “Mutation” clusters (labeled as 0–21, Figure S1A). Compared to gene isoform expression, somatic single nucleotide variant (SNV) profiles showed much diminished power of discrimination between cancer types, exemplified by the mixed composition of almost all the 22 clusters (Figure S1A-B). The cluster residence map (Figure 1F, left) better illustrates this observation with the nontrivial overlap between one Mutation cluster and multiple Isoform clusters, especially Mutation clusters 0-3. However, we found strong concordance between our isoform expression-based clustering results and the published multiomic features-based clustering (named “iCluster”) results [21] (Figure 1F, right). The iCluster scheme incorporates TCGA mRNA-seq, microRNA (miRNA)-seq, DNA methylation, reverse-phase protein array (RPPA), and DNA copy number data to partition all the TCGA tumor samples into 28 clusters (Figure S1C). Comparing to other TCGA-based clustering schemes with variation of information (VoI) analysis (see Methods), we observed the strongest concordance of our Isoform clustering scheme with cancer type and mRNA, followed by miRNA and iCluster (Figure 1G). The VoI analysis further shows that the gene isoform expression provides higher resolution of cancer type classification compared to mRNA at the gene level (1.6 vs. 2.1 in VoI), which is consistent with our previous finding that the differential isoform ratio has little correlation with the differential expression at gene level in the human kinome in SKCM cancer [4].

Taken together, both gene isoform expression and gene somatic mutation profiles provide unique molecular taxonomy in addition to the current organ/tissue histology-based pathology classification and extend beyond other phenotypic characteristics such as tumor stage and tumor tissue purity (Figure S1D). While isoform and mutation molecular signatures stratify cancer samples at different resolution, we propose that integration of their information content might better characterize the intrinsic molecular traits of tumor samples. Therefore, the sample-matched RNA-seq and DNA-seq data across multiple cancer types represent a great opportunity to investigate the association between gene somatic mutation and gene isoform expression patterns.

### SoMAS identifies somatic mutations associated with alternative splicing in an effective and efficient way

Based on the matched TCGA multi-omics data as described above, we developed the SoMAS pipeline to investigate the potential impact of gene somatic mutation on gene isoform expression (Figure 2A). The SoMAS workflow consists of two steps: In the first step, the high-dimensional gene isoform expression matrix (d=59,866 derived from the 15,448 multi-isoform genes) for each cancer type is trimmed by mean.var.plot method (built in the Seurat toolkit [22], see Methods) into a more informative expression matrix, which keeps only the most variable isoforms but still maintains high-dimensionality (d≈3,500). After that, the Principal Component Analysis (PCA) [23] is performed to further reduce the dimension of the informative expression matrix into a much lower-dimensional PC score matrix (d=50) by calculating a PC loading matrix [24]. Each column of the PC loading matrix performs a particular linear combination of the top variable isoforms into a meta-isoform, with the combination coefficients stored in the corresponding column of the PC loading matrix. All the meta-isoforms comprise the coordinates of the new low-dimensional expression space.

**Figure 2.**
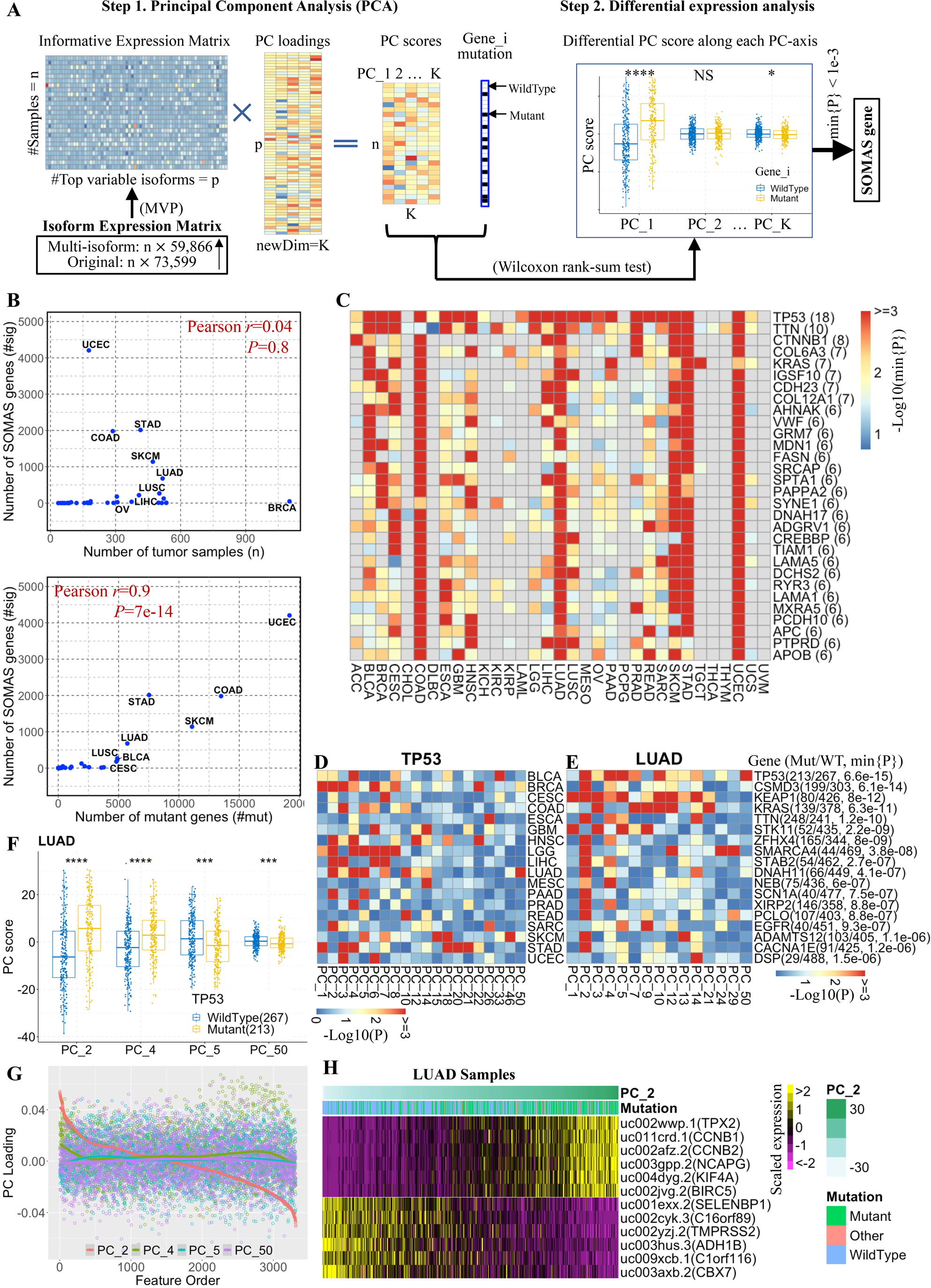
Overview of the SoMAS workflow and SoMAS genes across pan-cancer. (A) The SoMAS workflow consists of two steps: dimension reduction with PCA followed by a differential PC score analysis with Wilcoxon’s rank-sum test. SoMAS genes are determined if the minimum of p-values along all PC-coordinates is smaller than a predefined threshold. The MVP method (developed in Seurat) was used to trim the original isoform expression matrix into a more informative expression matrix which keeps the top variable isoforms only. (B) The number of SoMAS genes detected in each cancer against the sample size (upper) and the number of mutant genes (lower). (C) Heatmap shows the Log10-tranformed minimum p-value along the 50 PC coordinates for the top 30 SoMAS genes across pan-cancer. Genes were ranked by the number of cancer types in which they were detected as SoMAS genes, as indicated in parentheses. (D) TP53 was detected as a SoMAS gene in 18 cancer types, involving 20 PC axes as indicated. (E) The top 18 SoMAS genes detected in LUAD cancer. Heatmap shows the significance level of each gene across the 50 PC-axes. Number of wildtype and mutant samples, and the minimum p-value among the 50 PCs for each SoMAS gene are indicated. PCs not significant in any of the 18 genes in LUAD were excluded for clarity. (F) The details of differential PC score analysis for TP53 in LUAD cancer. TP53 was tested significant along four PC axes: PC_2, PC_4, PC_5 and PC_50 in LUAD. \*\*\*\**P*<1e-4, \*\*\**P*<1e-3 by Wilcoxon rank-sum test. (G) Visualization of the PC loading scores along significant PC axes for TP53 in LUAD cancer. Isoform features (x-axis) were ordered by PC loadings along PC_2 axis. (H) Scaled expression levels of the top 6 positive (rows 1-6) and top 6 negative (rows 7-12) isoforms of all samples ordered by their PC scores along PC_2 axis in LUAD cancer. The isoforms were ranked by the PC loading value along PC_2 axis (Figure 2A). See also Table S2 and Figure S2.

In the second step, a differential PC score analysis is conducted along each of the 50 PC-coordinates based on the mutation status (SNV) of the studied gene by Wilcoxon rank-sum test. Following the differential PC score analysis, the significant genes (termed SoMAS genes) are determined if the minimum of the 50 p-values (along the 50 PC coordinates) is smaller than a predefined threshold (*P*_min_<1e-3). It should be noted that, it is possible that one gene is tested significant along multiple PC coordinates (termed significant PCs hereafter). With these significant PCs we can trace back to the PC loading matrix to determine all transcripts (including both *cis*- and *trans*-positions) that are significantly associated with this SNV at the same time, and hence generate the association between this SNV and the abundance of those significant transcripts.

We applied SoMAS to 33 individual TCGA cancer types and detected a varied number of SoMAS genes (Table 1) among the different cancers. We identified more than 2,000 SoMAS genes in STAD and UCEC cancers which are known for hypermutability, but none in CHOL and KICH cancers. We obtained only one SoMAS gene in three cancer types: DLBC (*PIM1*), LAML (*RUNX1*) and UCS (*FBXW7*). Some genes were ranked in the top 3 most significant SoMAS genes (ranked by *P*_min_) in multiple cancer types, such as *TP53* (in 12 cancers) and the RAS family including *KRAS*, *HRAS* and *NRAS* (in 5 cancers). The number of SoMAS genes in each cancer type showed little correlation with the sample size (Pearson r=0.04, *P*=0.8), but showed significant correlation with the number of mutant genes detected in that cancer (Pearson r=0.9, *P*=7e-14), especially for cancers with more than 5,000 mutant genes (Figure 2B). We pooled these genes to create a master list of 7,140 unique (i.e., nonduplicated) SoMAS genes and ranked them by the number of cancer types in which they were detected as a SoMAS gene (Table S2A). These unique SoMAS genes were present in a varied number of cancer types (ranging from 1 to 18). Specifically, 2,558 out of the 7,140 genes (36%) were detected as SoMAS genes in two or more cancer types and the remaining 4,582 genes were significant in only one cancer type. Many well-known oncogenes (e.g., *CTNNB1* and the RAS family), tumor suppressors (e.g., *TP53* and *PTEN*) and RNA-binding protein genes (e.g., *RBM10*, *TNRC6A* and *SF3B1*) were detected as SoMAS genes in multiple cancers.

**Table 1.**
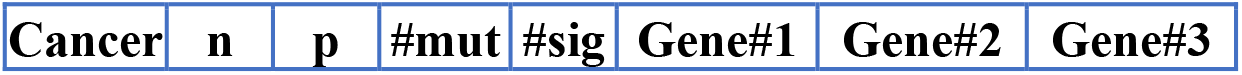
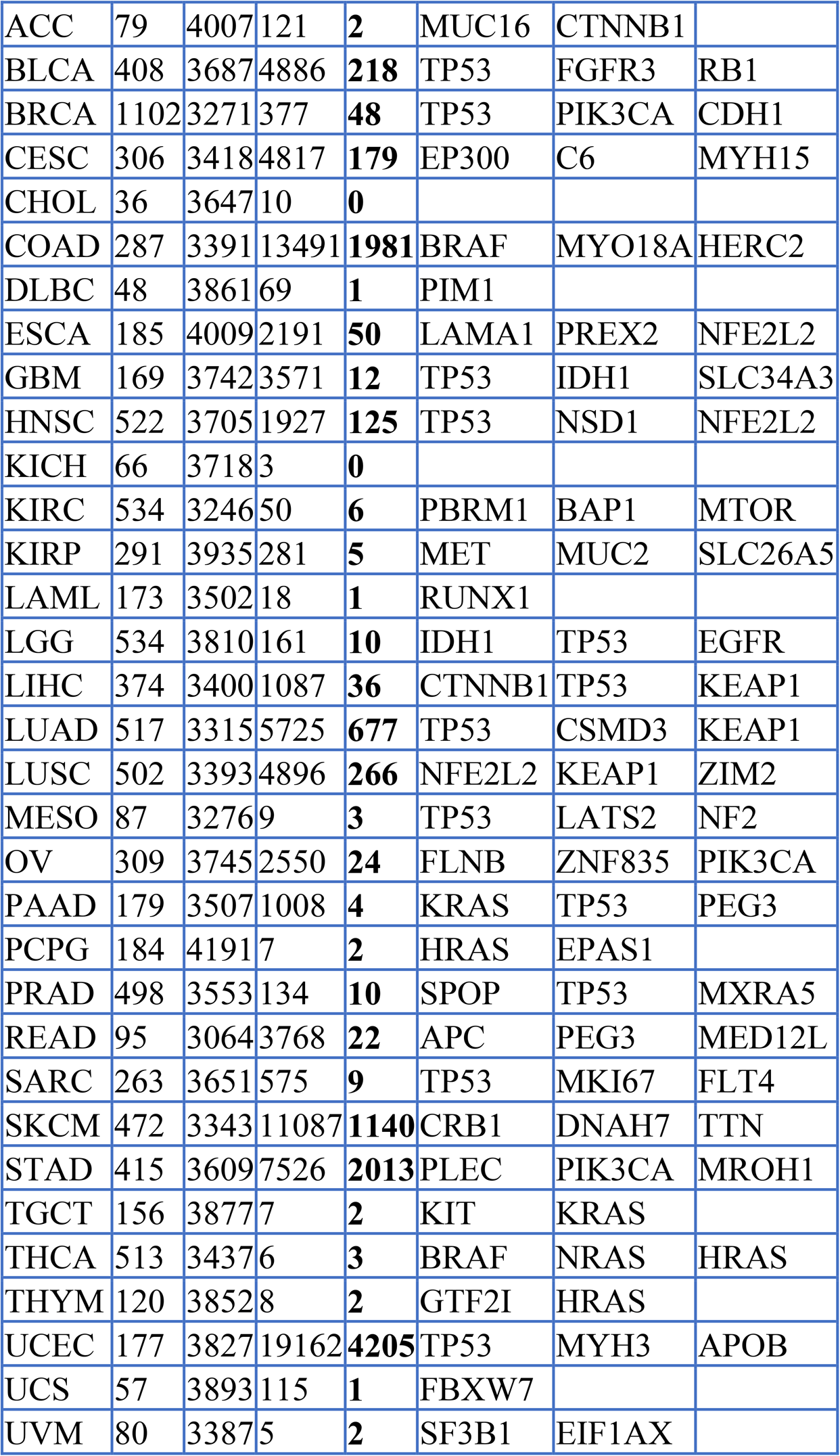
A summary of the SoMAS analysis for 33 TCGA cancer types. n: number of samples; p: number of top variable isoforms; #mut: number of genes mutated in ≥2% (and ≥5) samples; #sig: number of SoMAS genes detected; Gene#1-3: the top 3 SoMAS genes detected in each cancer type (ranked by the minimum p-value in the SoMAS analysis).

To assess whether disease-relevant genes were returned in our analysis, we examined overlap with The Cancer Gene Census (CGC) genes [25] curated in the latest COSMIC (Catalogue of Somatic Mutation in Cancer) database. Specifically, we found 102 out of the 733 CGC genes (14%) were present in the SoMAS gene list covering at least three cancer types (hypergeometric test P=1.1e-27), and this number increased to 411 (56%) overlapping all the 7,140 SoMAS genes (P=4.9e-41) (Table S2A).

Each of the top 30 SoMAS genes was significant in six or more cancer types, and 17 out of the 30 genes were shared by the top four cancer types (UCEC, STAD, COAD and SKCM) having the most SoMAS genes (Figure 2C, Table S2A). *TP53* turned out to be the most prevalent SoMAS gene, present in 18 out of 33 TCGA cancer types (Figure 2D).

As the SoMAS method utilizes PCA, we examined the contribution of different PC-axes to biological relevance. We observed that the SoMAS genes could be tested significant along any of the 50 PC-axes, demonstrated by the nontrivial percentages represented by the last 10 PCs (PC_41 – PC_50) (Figure S2A; Figure 2D, E). Some genes tested significant along multiple PCs in some cancers. This finding indicates that all the 50 dimensions in the new meta-isoform expression matrix (the PC score matrix) are essential for capturing the critical information contained in the original high-dimensional isoform expression matrix (with 59,866 isoforms derived from 15,448 multi-isoform genes, see Figure 2A). For example, *TP53* tested significant along four PC axes in the differential PC score analysis step, in the sense that *TP53* mutation was associated with increased PC score along PC_2 and PC_4 but decreased PC score along PC_5 and PC_50 (Figure 2F). We noticed that the PC loading values along different PC-axes are quite different among each other (Figure 2G), which is sensible in that PCA tries to capture complementary information inherent in the input matrix through different coordinates in the transformed space.

We mathematically proved that the significant association between a gene mutation and a PC score is equivalent to the significant association between the gene mutation and the expression level of isoforms with large (positive or negative) PC loading values, owing to the linear combination property involved in the PCA transformation (Methods). This equivalence can be better illustrated by the *TP53* example in the LUAD cancer (Figure 2H; Figure S2B). Specifically, the scaled expression level of the isoforms with high PC loading weights turned out to be well correlated (in positive or negative direction) with the PC score along the same PC-coordinate. Incrementally pooling the top target isoforms together well approximates the PC score along the significant PC axis, as manifested by the same example of *TP53* in LUAD (Figure S2C). With direct differential isoform expression analysis, we verified that *TP53* mutation was indeed significantly associated with the expression level of most (98%) of the top 100 isoforms (50 positive and 50 negative) (Figure S2D). However, the gene mutation can be associated with different isoforms of a gene at different significance levels, or even significantly associated only with some particular isoform(s) of that gene (Figure S2E).

We further confirmed that the mutation status of some SoMAS genes is significantly associated with the expression of their target genes in dual directions. Specifically, a SoMAS gene mutation can significantly (Wilcoxon’s P<0.05) upregulate some isoforms while downregulate the other isoforms of the same target gene. In total, we found 4,986 (i.e., 69.8% of all 7,140) SoMAS genes exhibited dual-direction associations with at least one (pooled mean: 2; range: 1-68) gene representing their top 100 target isoforms (Figure S2F-G, Table S2B). Interestingly, *TP53* showed highest significance level in dual-direction association in six cancer types (Figure S2H). These data imply that, besides impacting the expression level of isoforms individually, the SoMAS gene mutation can also affect the splicing process of their target genes, manifested by the upregulation and downregulation of particular isoforms of the same targe gene.

These analysis results clearly demonstrate that our proposed SoMAS pipeline can effectively identify all gene isoforms whose expression level is significantly associated with the mutation status of a designated gene simultaneously, which is much more efficient than the traditional differential expression analysis applied directly on the original isoform expression. We achieved this by employing the well-established dimension reduction technique PCA to transform the high-dimensional isoform expression matrix into a much lower-dimensional PC score matrix, followed by a differential PC score analysis instead of differential isoform expression analysis which involves a huge number of mutation-isoform pairs. Next, we will take a closer look at the SoMAS genes and their associated gene isoforms (transcripts) having the largest PC loading weights determined in each cancer type, to explore their biological and clinical meaning. Those top (primarily top 100 unless otherwise stated) isoforms significantly associated with a SoMAS gene are termed the target isoforms (direct or indirect) of this SoMAS gene in the subsequent analysis.

### SoMAS genes tend to associate with gene isoform expression in a *trans*-regulation manner

To further explore the SoMAS genes and their associated gene isoforms (targets) derived in each cancer, we first examined the genomic distribution of the top isoform targets of the SoMAS genes in each cancer type. The top 24 isoform targets of a SoMAS gene are typically distributed across 10-15 chromosomes in any cancer type (in all cancers pooled, median = 12). In all 31 TCGA cancer types with SoMAS genes detected, the top 100 isoform targets of each SoMAS gene typically covered 20 or more chromosomes (pooled median = 21) (Figure 3A). This observation indicates that a SoMAS gene can potentially impact the gene isoform expression in a genome-wide sense.

**Figure 3.**
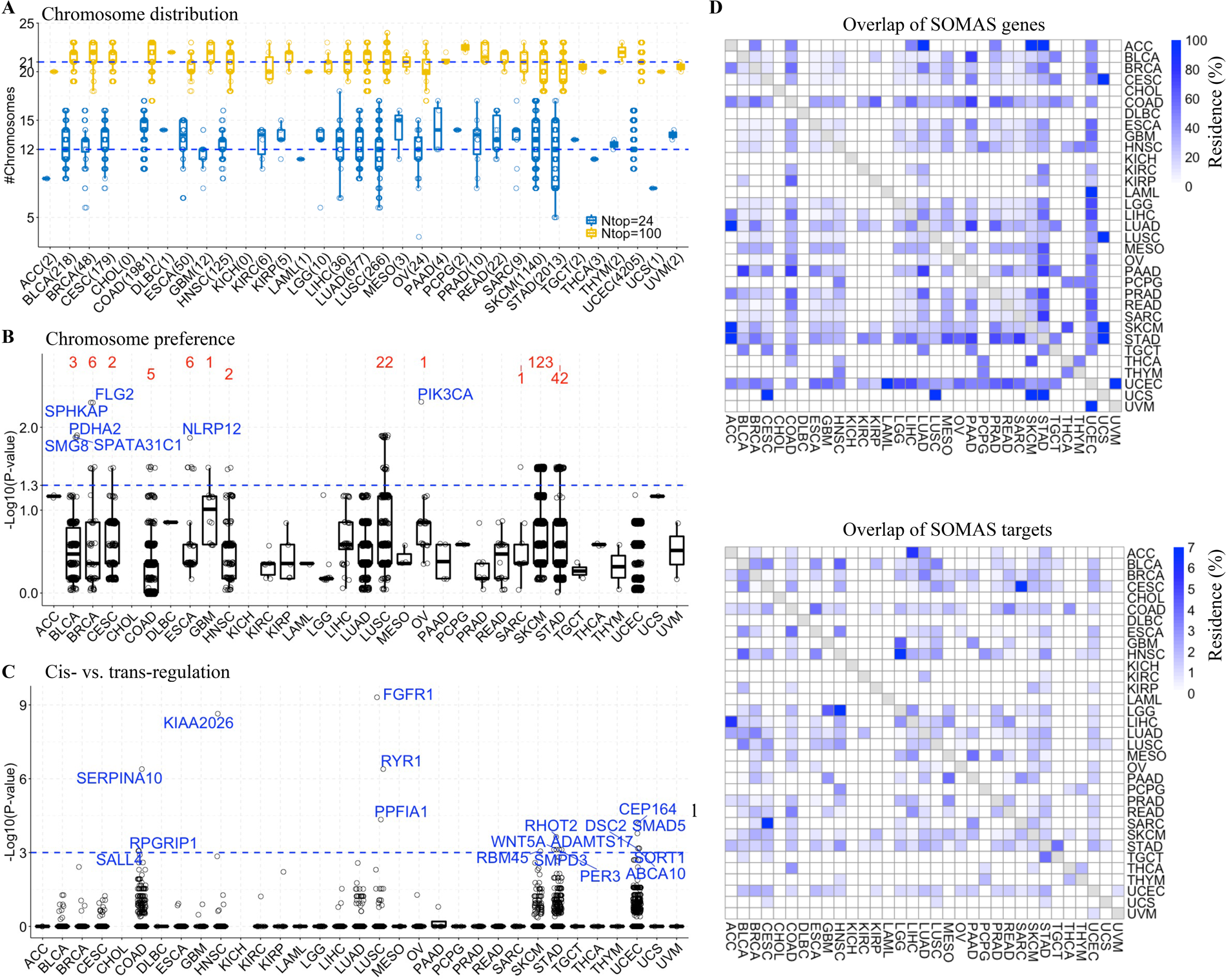
*Cis*- and *trans*-regulation patterns of SoMAS genes across pan-cancer. (A) Chromosome distribution of target isoforms of the SoMAS genes detected in each cancer type. Y-axis represents the number of chromosomes across which the top 24 and 100 target isoforms are distributed. Dotted lines indicate the pooled median for all cancer types. Numbers in parentheses in the x-axis labels indicate the number of SoMAS genes detected in each cancer (Table 1). (B) Chromosome preference of target isoforms of the SoMAS genes. For each SoMAS gene detected in each cancer type, the frequency distribution of its top 100 target isoforms along 24 chromosomes was compared to the number distribution of all isoforms across the 24 chromosomes by KS test. Y-axis represents the Log10-transformed p-value derived from the KS test. The number above each box whisker indicates the number of SoMAS genes yielding P<0.05 in the KS test in corresponding cancer type. (C) Boxplot illustrates the likelihood that SoMAS genes target isoforms in a *cis*-acting mechanism. Y-axis represents the Log10-transformed p-value derived from the hypergeometric test measuring the likelihood that the top 100 target isoforms of a SoMAS gene are in its *cis*-regulation region (±1Mb of TSS). Numbers of SoMAS genes with P<0.001 in the hypergeometric test and the most significant SoMAS gene are indicated above corresponding cancer types. (D) Overlap of SoMAS genes (upper) and SoMAS targets (lower) among cancer types. The top 100 isoform targets of each SoMAS gene were extracted and pooled together for each cancer type, followed by a pair-wised overlap calculation for all 33 cancer types. The percent of overlap (Residence) between two SoMAS gene sets was calculated as the size of overlap divided by the size of the smaller set. See also Table S3 and Figure S3.

Next, we checked whether there is any chromosome preference for the isoform targets of a SoMAS gene in each cancer type. To do this, for each SoMAS gene detected in each cancer type, we compared the frequency distribution of the top 100 isoform targets along the 24 chromosomes to the number distribution of all isoforms (Niso) across the 24 chromosomes and calculated a p-value for this comparison with the Kolmogorov–Smirnov (KS) test. We observed few SoMAS genes that passed the KS test at the P=0.05 (or −Log10(P)=1.3) level (Figure 3B; Figure S3A), i.e., showing significantly different probability distributions. This means that, the isoform targets show little chromosome preference for most SoMAS genes in any cancer type.

Now that SoMAS genes are associated with isoform expression of many genes across the whole genome without significant chromosome preference, we further ask whether SoMAS genes target their neighboring genes (or *cis*-region, defined as ±1Mb of its TSS) more often than distal genes (*trans*-region, defined as non-*cis*-regions). To address this question, we conducted a hypergeometric test to measure the significance level that a particular number (or more) of the top 100 isoform targets were located in the *cis*-region of a designated SoMAS gene (Figure S3B). We observed that very few SoMAS genes passed the hypergeometric test as a *cis*-regulation SoMAS gene in all cancer types at the *P*=0.001 level (Figure 3C). The number of *cis*-regulation SoMAS genes increased a little at the *P*=0.05 and *P*=0.01 levels but was still small (typically less than 5%). While it’s hard to claim that all the remaining genes are *trans*-regulation SoMAS genes, the SoMAS genes with a hypergeometric test *P*>0.999 (which means the test with the opposite null hypothesis gets a p-value of ∼0.001) took the majority (Figure S3C, Table S3). These results indicate that only a tiny part of SoMAS genes detected in each cancer type assumedly act through *cis*-regulation mechanism, while the majority are potential *trans*-regulatory SoMAS genes.

While most (64.2%) of SoMAS genes are cancer type-specific (Table S2A), i.e., an individual gene is detected as a SoMAS gene in only one cancer type, we found that the whole SoMAS gene set of many cancer types largely overlapped (see Method) with that of other cancer types (Figure 3D, upper). The SoMAS gene sets of 10 cancers, including UCEC, COAD, STAD, LUAD, SKCM, HNSC, CESC, BLCA, BRCA and LIHC, overlapped with that of 20 or more cancer types. Even for those cancer types with only one SoMAS gene detected, the single SoMAS gene was likely shared by multiple cancer types. This includes LAML (*RUNX1* was shared by 2 cancer types and UCS (*FBXW7* was shared by 4 cancer types). The only exception is DLBC, whose single SoMAS gene *PIM1* was specific to this cancer type. Interestingly, these inter-cancer overlapping patterns of SoMAS genes extended to show overlap among the isoform targets of those SoMAS genes detected among all cancer types studied (Figure 3D, lower). These results demonstrate an underlying commonality in the regulation of at least some alternative splicing events associated with somatic mutations across various cancer types.

Looking into the specific [SoMAS gene]:[SoMAS target] pairs, we observed distinct patterns in inter-cancer overlap profiles compared to that with SoMAS genes or SoMAS targets alone. While apparent pair-wise overlap in [SoMAS gene]:[SoMAS target] pairs among cancer types was still observed, most pairs turned out to be cancer type-specific (Table S3). Specifically, only 9,668 out of the 1,096,083 (0.9%) pooled unique [SoMAS gene]:[SoMAS target isoform] pairs are present in two or more cancer types; and this ratio increased slightly to 1.3% (12,997 out of 988,781) for pooled unique [SoMAS gene]:[SoMAS target gene] pairs. In both cases, we considered the top 100 target isoforms (and their corresponding genes) of each SoMAS gene. No [SoMAS gene]:[SoMAS target isoform] pairs or [SoMAS gene]:[SoMAS target gene] pairs were present in more than four cancer types. This means that a SoMAS gene tends to target different genes in different cancer types.

### SoMAS genes are significantly involved in tumor growth and metastasis related processes

We have detected a series of both common and cancer type-specific SoMAS genes in human cancers and verified their significant association with expression of multiple gene isoforms. We also showed that SoMAS genes typically associate with gene isoform expression across the whole genome, without obvious chromosome preference. Here we ask whether these SoMAS genes have any biological and/or clinical meaning to human oncogenesis. To address this question, we performed a series of biological process enrichment analyses and survival analyses to investigate the biological and clinical relevance of the SoMAS genes, respectively.

We conducted a KEGG signaling pathway enrichment analysis on the top 908 “pan-cancer” SoMAS genes that cover at least three cancer types (Table S2A) to study their biological functions. This analysis revealed that these 908 genes are significantly enriched in the cell migration (e.g., Focal adhesion, ECM-receptor interaction, Protein digestion and absorption) and cell proliferation (including Growth hormone synthesis/secretion/action and PI3K-Akt signaling pathway) related signaling pathways (Figure 4A, left; Table S4). Similarly, we checked the KEGG pathway enrichment profiles of the top genes ranked by the SoMAS pipeline (including those not meeting the original significance threshold p <1e-3 set for a SoMAS gene) in each individual cancer. Due to the dramatically varied number of SoMAS genes detected in different cancer types (Table 1) and for consistency with the pan-cancer enrichment analysis, we also considered the top 908 (if available) mutant genes that obtained a P_min_<0.05 (instead of P_min_<0.001 for a SoMAS gene) in the SoMAS analysis (Step 2 in Figure 2A) in the cancer-specific enrichment analysis. In this way, the same top 30 KEGG pathways turned out to be significant in multiple cancer types in the cancer-specific enrichment analysis (Figure 4A, right). While a large part (n=610; 67%, Table S2A) of the 908 pan-cancer SoMAS genes only covered three cancer types, a large fraction (n=22; 73%, Figure 4A, right) of the top 30 significant pan-cancer pathways proved significant in more than 3 cancer types. This implies that different member genes of the same pathway were detected as SoMAS genes in different cancers.

**Figure 4.**
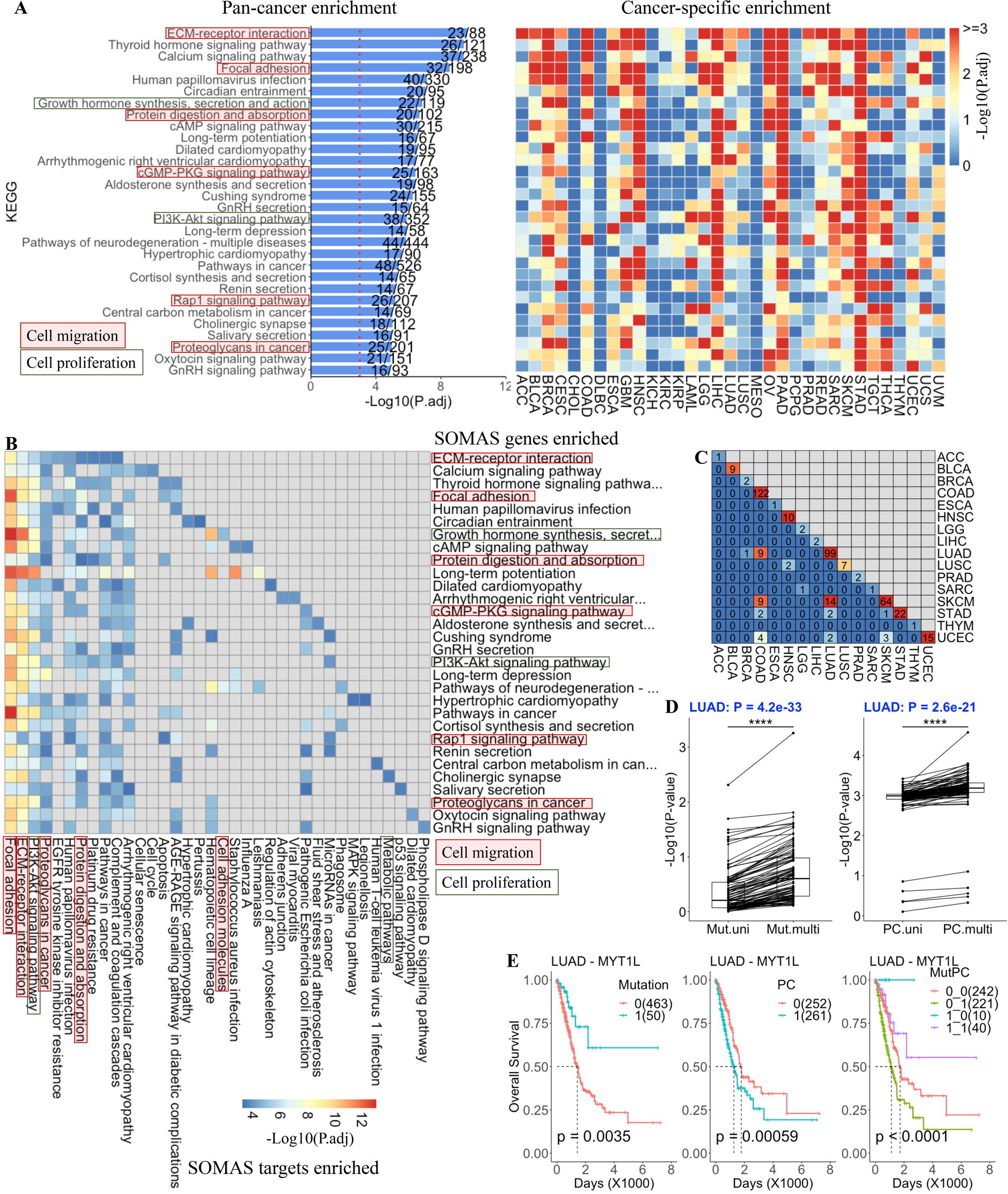
Biological and clinical significance of the SoMAS genes. (A) Signaling pathway enrichment analysis on the top 908 SoMAS genes (from 22,029 background genes) that are tested significant in ≥3 cancer types against 343 KEGG pathways in both pan-cancer (bar plot, left) and cancer-specific (heatmap, right) manners. Only top 30 enriched pathways are shown. In the cancer-specific enrichment analyses, gene numbers were trimmed to the top 908 (still subject to *P*<0.05 in the SoMAS analysis) for a fair comparison. Number of genes contained in each pathway (n) and number of significant genes hitting that pathway (m) are shown on each bar (m/n). Cell migration and proliferation related pathways are marked. (B) KEGG enrichment analysis on the target isoform genes of SoMAS genes hitting each of the top 30 pathways shown in (A). For each pathway, the top 100 target isoform genes of each SoMAS gene hitting that pathway were picked and pooled (with duplicates removed) to perform the enrichment analysis. Gene numbers were trimmed to the top 908 based on the frequency of presence of each gene in the pooled list. Only top 10 KEGG pathways enriched by the SoMAS targets were used to generate the final heatmap. (C) Pairwise overlap of Additive SoMAS genes detected across 16 cancer types. The other 17 cancer types without Additive SoMAS genes detected are ignored. (D) Paired *t*-test compares the p-values between univariate and multivariate regression analysis for mutation (left) and its corresponding PC score (right) of the Additive SoMAS genes in LUAD cancer. (E) Survival analysis based on top Additive SoMAS gene MYT1L in LUAD cancer. LUAD samples were divided into groups based on the mutation status of the gene (left), binarized PC score (middle) and combination of them (right). Mutation: 0=wildtype, 1=mutant; PC: 0=negative, 1=positive PC score. MutPC: Combination Mutation_PC score. Sample size of each group is indicated. P-values were derived from log-rank test. See also Table S4 and Figure S4.

Similarly, we performed a gene set enrichment analysis on the top 908 pan-cancer SoMAS genes against 50 well-known Hallmark gene sets collected in the Molecular Signature Database (MSigDB) [26]. Consistent with the KEGG pathway enrichment profiles, the top 908 SoMAS genes showed significant enrichment in cell migration and cell proliferation related gene sets in both pan-cancer and cancer-specific manners (Figure S4A; Table S4). In particular, the top 5 enriched gene sets, including cell migration related Epithelial_Mesenchymal_Transition and cell proliferation related Mitotic_Spindle gene sets, covered 13-17 cancer types. This further confirmed that different member genes of the same gene set exert their association to isoform regulation in different cancers.

We further examined the biological function of the target isoforms of the top 908 pan-cancer SoMAS genes by KEGG pathway enrichment analysis. The most striking observation is that these SoMAS target genes are significantly enriched in nearly the same cell migration and cell proliferation related signaling pathways as the SoMAS genes (Figure 4B). Besides, the SoMAS targets are enriched in other oncogenic signaling cascades that assumedly represent a critical complement to the SoMAS genes. Considering the observation that SoMAS genes tend to regulate gene isoform expression in a *trans*-acting manner, these results imply that SoMAS gene mutations and associated isoform expression alternations might work in a synergistic way to regulate human cancer progression.

To test this synergy hypothesis, we conducted both univariate and multivariate Cox regression analysis on the two associated variables in all tumor samples of a cancer type: the SoMAS gene mutation status and its associated PC score (along the significant PC of this SoMAS gene) (see Methods). We identified a proportion of SoMAS genes with enhanced power in distinguishing survival groups when integrated with their associated PC scores in 16 cancer types (Figure S4B), indicating a synergetic or additive effect. The additive effect refers to the higher significance level (or smaller p-value) obtained with the combination of the two variables compared to that with each variable alone in the Cox regression model. For simplicity, hereafter, we termed these special SoMAS genes as additive SoMAS genes (Table S4). We observed generally negligible overlap of additive SoMAS genes among cancer types (Figure 4C). Furthermore, direct log-rank test confirmed this additive effect between gene mutation and PC score (binarized as 0/1 for simplicity) in representative cancer types LUAD (Figure 4D-E) and BLCA (Figure S4C-D). Noting that the PC score is a linear combination of multiple gene isoforms, this synergistic effect further indicates a functional outcome uniting gene mutation and gene isoform expressions for some SoMAS genes in particular cancer types.

### RBP, SF and TF genes are largely detected as SoMAS genes in multiple cancer types

RNA-binding proteins (RBPs) and splicing factors (SFs) are known to play a vital and direct role in shaping the gene isoform expression landscape [27]. On the other hand, the expression of RBP and SF genes themselves are tightly regulated by a network of transcription factors (TFs). In this sense, the TFs are also able to impact gene isoform expression in certain way and theoretically can be detected as SoMAS genes by our system. Therefore, we asked what fraction of RBP, SF and TF genes can be identified as SoMAS genes by SoMAS. To address this question, we curated 1,681 RBP genes from MSigDB [26], 394 SF genes from published literature [16] and 2,478 TF genes from public resource (Methods), which had to overlap genes within the somatic mutation data from TCGA (Table S5).

Generally, our pipeline detected a considerate number of RBP, SF and TF genes as SoMAS genes present in one or more cancer types (Figure 5A). Particularly, a significant part of RNA-binding SF genes (mathematically denoted RBP∩SF) were detected as SoMAS genes in at least one cancer type (RBP ∩ SF ∩ SoMAS, Hypergeometric test P=0.032); whereas the overlap between SoMAS and RBP (RBP∩SoMAS) or SF (SF∩SoMAS) genes alone turned out to be not significant. We observed 10 RNA-binding SFs were detected as SoMAS genes in three or more cancer types, including four DEAD box protein genes (DDX18, DHX34/40/57). The DEAD box protein genes encode RNA helicases that participate in several important cellular processes including spliceosome assembly [28]. These data imply that our SoMAS approach is more likely to identify RNA-binding SFs as splicing-relevant factors (i.e., SoMAS genes) compared to general RBPs or SFs.

**Figure 5.**
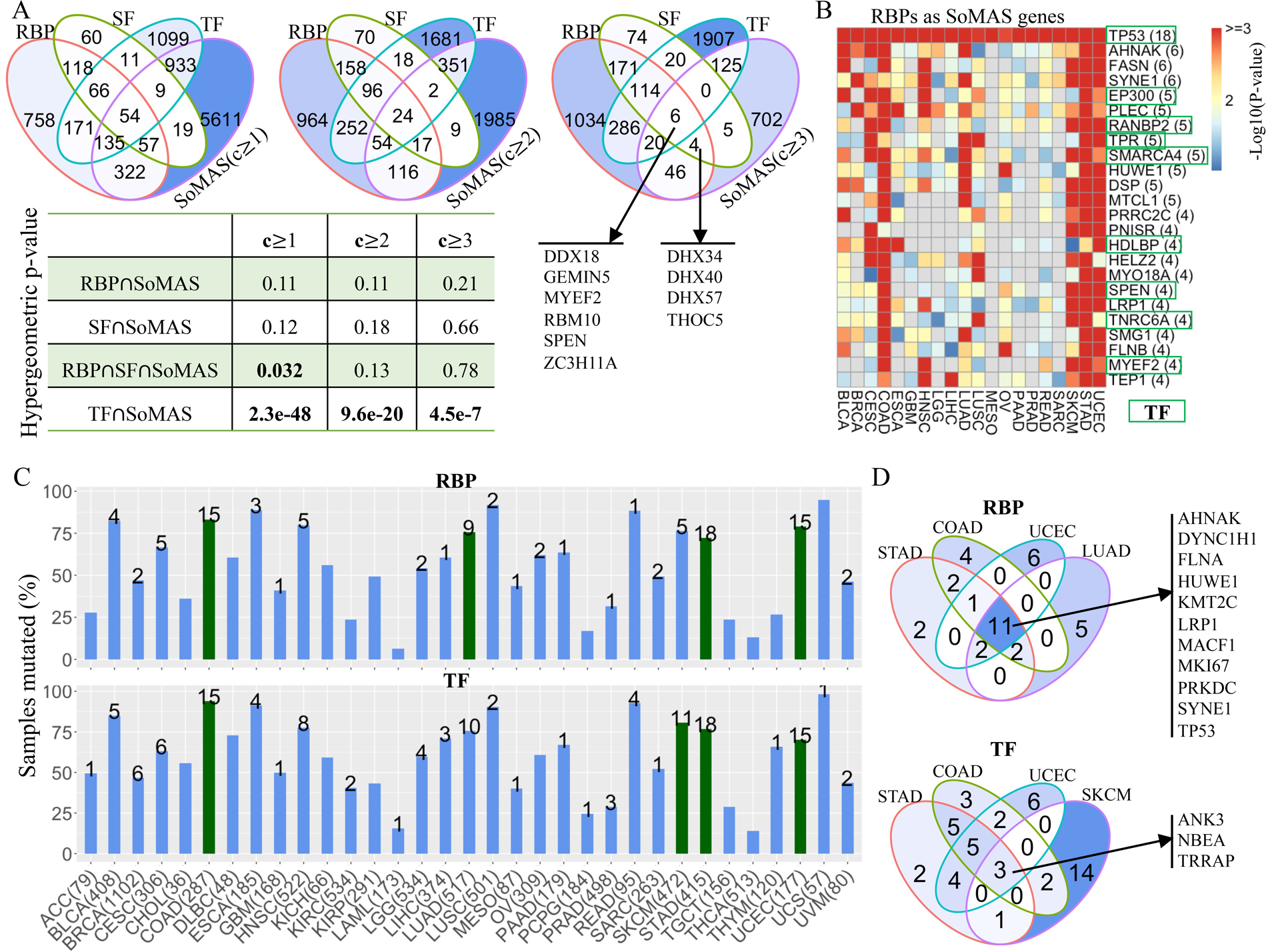
RNA-binding protein (RBP), splicing factor (SF) and transcription factor (TF) genes detected as SoMAS genes and their mutational landscape across cancer types. (A) Venn diagram shows overlap among RNA-binding protein (RBP), splicing factor (SF), transcription factor (TF) and SoMAS genes with increasing pan-level of cancer coverage c≥1, 2, 3. P-values were derived from hypergeometric test, with 22,029 mutant genes as background. (B) Heatmap shows the Log10-tranformed p-values of the 24 RBP genes detected as SoMAS genes in ≥4 cancer types. Number of cancer types are indicated in parentheses. (C) Percent of tumor samples mutated in at least one of the 20 most frequently mutated RBP (upper) and TF (lower) genes in each cancer type. The number above a bar indicates the number out of the 20 RBP/TF genes detected as SoMAS genes in the corresponding cancer type. The number of patients with gene isoform expression data available in each cancer type is also provided. The top four cancer types with most (out of the top 20) RBP and TF genes detected as SoMAS genes are colored green. (D) Overlap of top 20 RBP (upper) or TF (lower) genes among four cancer types. See also Table S5 and Figure S5.

Interestingly, the TFs were widely detected as SoMAS genes at all pan-levels (i.e., covering at least 1-3 cancer types, respectively, Figure 5A), including some RBP intersected TF genes (see RBP∩SF∩SoMAS in the Venn diagrams). Particularly, among the 24 RBP genes that were detected as SoMAS genes in ≥ 4 cancer types, nine are TF genes, including the cell cycle regulators TP53 and EP300, and the chromatin modifiers SMARCA4 and SPEN (Figure 5B, Table S5). Notably, six of the 10 RNA-binding SFs are also TFs.

We also checked the fraction of RBP, SF and TF genes serving as isoform targets of our detected SoMAS genes. We observed that 34% (572/1681) of RBP genes, 32% (127/394) of SF genes and 44% (1,079/2,478) of TF genes ranked in the top 100 target isoforms of some SoMAS genes in some cancer types (Table S5). For example, the splicing factor HNRNPK gene (specifically, one or more HNRNPK isoforms) represents one of the top 100 target isoforms of a varied number of SoMAS genes across 11 cancer types, from one SoMAS gene in CESC, GBM, HNSC, PCPG and READ to 21 SoMAS genes in STAD. These results imply that, besides direct regulation of downstream isoform splicing/transcription by individual SoMAS genes, isoforms can be impacted by two alternative scenarios: (i) multiple SoMAS genes can target the same gene for splicing and (ii) SoMAS genes can exert indirect effects by directly affecting splicing factors, which then regulate the splicing of other genes.

Consistent with their importance to gene splicing and transcription regulation, respectively, the RBP and TF genes were found to be highly frequently mutated in human cancer samples in general (Figure 5C). Specifically, a median of 56% and 60% of tumor samples bear somatic mutations in at least one of the top 20 (by mutation frequency) mutated RBP and TF genes across the 33 TCGA cancer types studied, respectively. In addition, out of the 151 TF genes that were detected as SoMAS genes in at least three cancer types (see the third Venn diagram in Figure 5A), 67 genes have shared TF targets and SoMAS targets (Table S5). For example, 352 (27%) of predicted SoMAS targets of TP53 are also documented TF targets of TP53. This percentage increased to 41% for the tumor suppressor MZF1 [29] and 45% for proto-oncogene FLI1 [30], respectively. These results indicate that the SoMAS genes can regulate gene isoform expression by acting as TFs.

The most frequently mutated RBP and TF genes include many SoMAS genes, meaning they are associated with altered isoform expression. Specifically, we detected a varied number of SoMAS genes from the top 20 frequently mutated RBP genes in 21 cancer types (Figure 5C, upper). Eleven RBP genes were shared by the top four cancer types (STAD, COAD, UCEC and LUAD, ranked by the number of SoMAS genes detected from the top 20 frequently mutated RBPs) in the list of their own top 20 mutated RBPs (Figure 5D, upper). Similarly, we identified different numbers of SoMAS genes from the top 20 frequently mutated TF genes in 26 cancer types (Figure 5C, lower). Only three genes were shared by the top four cancer types (STAD, COAD, UCEC and SKCM) in the list of their own top 20 mutated TFs (Figure 5D, lower). These results indicate that: (i) both the top frequently mutated RBP and TF genes tend to enrich for SoMAS genes in human cancers and (ii) top mutant RBP genes are more likely to be shared by many cancer types compared to top mutant TF genes, and hence have less tissue specificity. Interestingly, in most samples of each above highlighted cancer types, the top 20 mutated RBP and TF genes are mutated in a nearly mutually exclusive manner (in the sense that each sample within a cancer type only bears mutations on one or very few of the 20 genes) (Figure S5). This exclusivity suggests a functional redundancy among the top 20 mutated RBP/TF genes in each cancer type [7], implying that they may converge to a same signaling pathway or a gene set of particular biological process [31].

Taking the above analysis results regarding RBF, SF and TF genes together, we conclude that SoMAS can detect many RBP, SF and TF genes that are critical for gene splicing/transcription regulation in both pan-cancer and cancer type-specific sense.

### Co-profile of chromatin accessibility around SoMAS genes and targets is largely disrupted

We have established that the mutation of a SoMAS gene can be significantly associated with the isoform expression pattern of up to hundreds of genes (called targets) across the whole genome. While the rationale and mechanisms underlying these associations are multifaceted, most of them involve the direct or indirect impact of somatic mutations on the splicing/transcription machinery interacting with the SoMAS gene itself or its target genes [32]. In this sense, we hypothesize that the chromatin accessibility status surrounding the SoMAS gene and its associated target genes may undergo certain changes corresponding to the potential shift in the DNA-protein and/or RNA-protein interactions. To address this hypothesis, we resorted to the sample-matched ATAC-seq data [33] in parallel with the TCGA DNA-seq and RNA-seq data used in our SoMAS pipeline. We examined the association between the chromatin accessibility profiles surrounding a SoMAS gene and the mutation status of this SoMAS gene. Meanwhile, we checked the correlation between the isoform expression of the associated SoMAS targets and their corresponding local chromatin accessibility (Methods). The analysis of association between somatic mutation and local chromatin accessibility covered a median of 15.9% of the SoMAS genes detected in 16 TCGA cancer types with available ATAC-seq data. Generally, the chromatin accessibility in the intron and promoter regions showed more significant association with the SoMAS gene mutation compared to the other four annotated regions including Distal, Exon, 3’UTR and 5’UTR (two-sided Wilcoxon rank-sum test on −Log10(HMP, or harmonic mean p-value), P=8.8e-20) (Figure 6A, upper; Figure 6B; Table S6). This tendency is even more discernible in the parallel study detecting correlation between isoform expression and chromatin accessibility (Wilcoxon P=4.0e-100) (Figure 6A, lower; Figure 6C; Table S6). These results indicate that both gene somatic mutation and gene isoform expression tend to have a significant association with the local chromatin accessibility status (at the level of HMP<0.05), and the significance level turned out to be genomic region-specific. It should be noted that these data are insufficient to determine the causative relationship among these three processes (i.e., somatic mutation, isoform expression and chromatin accessibility), which necessitates further experimental investigations.

**Figure 6.**
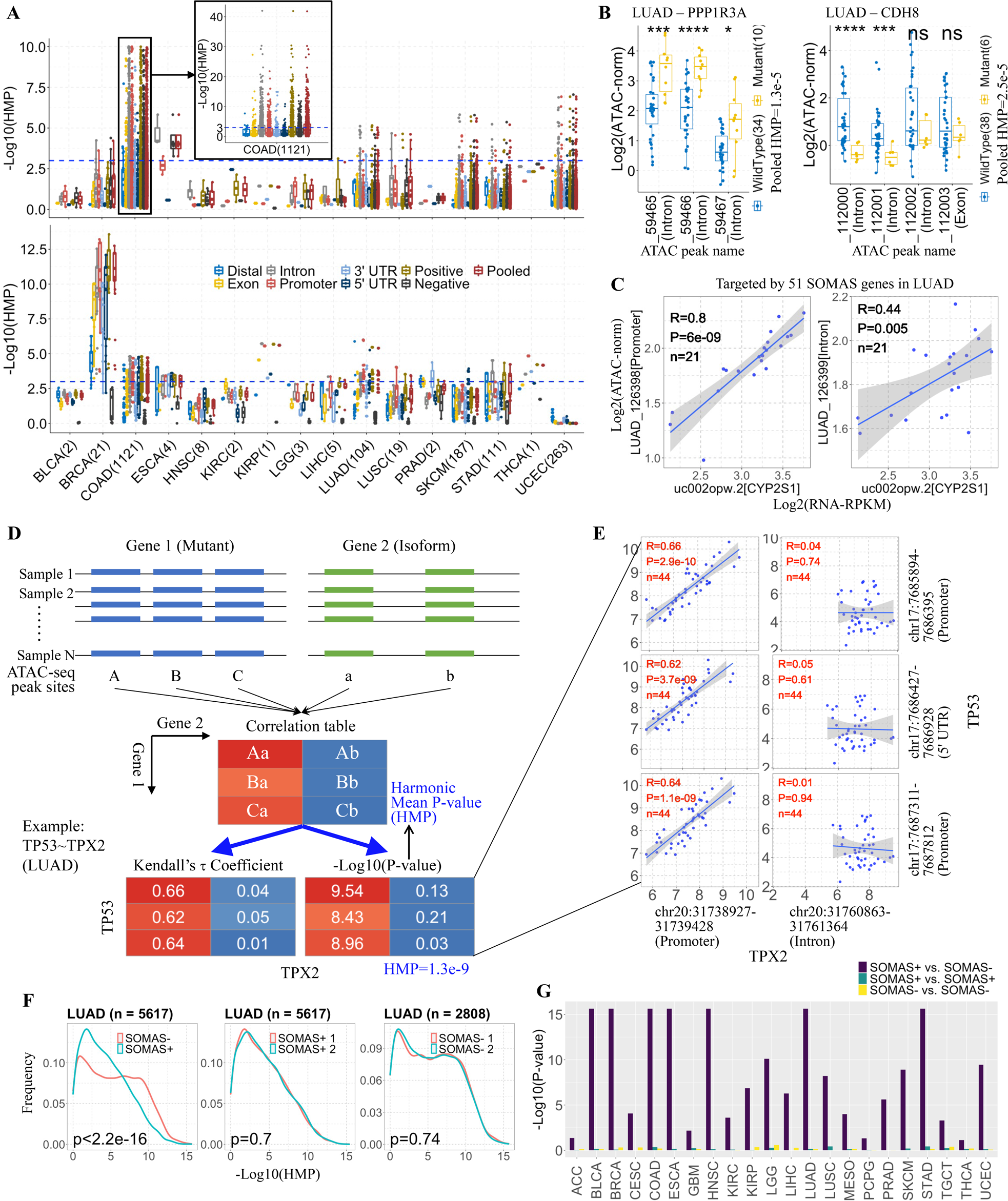
Chromatin accessibility status around SoMAS genes and targets. (A) Boxplots show the association between gene mutation (upper, by student *t*-test) or isoform expression (lower, by Kendall correlation analysis) and chromatin accessibility on SoMAS genes and targets in 16 cancer types. The number of SoMAS genes covered in the *t*-test in each cancer type was indicated in the x-axis (SoMAS genes without ATAC peak sites were skipped). The y-axis represents the Log10-transformed harmonic mean *p*-value (HMP) calculated for ATAC peak sites of each category as indicated. Top 100 target isoforms of each SoMAS gene were combined to calculate the HMP in the lower panel. Log10HMP level of 1.3 and 3 (corresponding to HMP level of 0.05 and 0.001, respectively) are marked by blue dashed line. Positive: ATAC-seq sites yielding a positive association between gene mutation (Fold change of mutant over wildtype > 0) or isoform expression (Kendall’s τ coefficient > 0) and chromatin accessibility; Negative: opposite to the Positive scenario; Pooled: HMP was calculated on all ATAC peak sites overlapping the gene set of interest. (B) Example SoMAS genes with significant association (*t*-test) between gene mutation and chromatin accessibility in LUAD cancer. Only pooled HMP was indicated for each gene. The x-axis indicates the names of peaks (given by the ATAC-seq data with cancer type prefix LUAD_ omitted) overlapping with the SoMAS gene in study. The y-axis Log2Norm(ATAC) value indicates the log2-transformed normalized ATAC-seq insertion counts. \*\*\*\**P*<1e-4, \*\*\**P*<1e-3, \**P*<0.05. (C) An example isoform targeted by multiple SoMAS genes with significant positive correlation (Kendall) between isoform abundance and chromatin accessibility in LUAD cancer. The x-axis refers to the isoform and corresponding gene name, and the y-axis refers to the name and genomic annotation of the ATAC peaks falling in the isoform locus. (D) Schematic of workflow for assessing the correlation in chromatin accessibility between SoMAS genes (mutant) and their targets (isoform). The correlation table lists correlation profile (including Kendall’s τ coefficients and p-values) of all possible combinations of ATAC-seq peak sites located in mutant and isoform genes. The HMP was calculated based on the p-values in the correlation table. (E) Detailed correlation status of a representative gene pair (TP53 vs. TPX2) in LUAD cancer. Shown are pairwise correlations between 3 ATAC peak sites on TP53 and 2 sites on TPX2. (F) Frequency distribution of Log10-HMP values (based on pairwise Kendall correlations) for gene pairs from both SoMAS (SoMAS+) and nonSoMAS (SoMAS-) pools (left), SoMAS+ only (middle) and SoMAS-only (right) in LUAD. The number of gene pairs for each scenario is shown above each panel. The p-value for significance level of distribution difference was derived from Kolmogorov-Smirnov (KS) test. (G) Summary of KS test p-values (Log10-transformed) for all 33 TCGA cancer types. The grouping strategy was same to (F). P-values < 2.2e-16 were trimmed to 2.2e-16 for better visualization. See also Table S6 and Figure S6.

Next, we checked the combinatorial profiles of chromatin accessibility by pairing the mutant gene (SoMAS gene) locus and associated target isoform (SoMAS target) locus with Kendall correlation analysis (Figure 6D). The correlation analysis was conducted in a pairwise manner between all ATAC peak sites located in the mutant gene against those located in the target isoform, which involves multiple combinations for genes with multiple ATAC peak sites. Figure 6E illustrates a specific example of the correlation analysis between TP53 and TPX2 in LUAD, which again displayed obvious bias specific to genomic region types (the correlation was significant in TPX2’s promoter region but not in its intron region).

Integrating all p-values obtained in the pairwise correlation analysis for each gene pair (SoMAS gene vs. SoMAS target), we calculated the HMP value to assess the overall correlation in chromatin accessibility between the two genes in the pair. To do so, we first calculated HMP for both SoMAS pairs (here a SoMAS pair refers to a SoMAS gene and one of its associated isoform targets) and non-SoMAS pairs (a randomly chosen pair from outside the SoMAS pairs pool). Then we checked the frequency distribution of the HMPs of the two scenarios. We found that the SoMAS pair group invariably exhibited a distinct HMP frequency distribution pattern compared to the non-SoMAS pair group with KS P<0.05 in all 33 TCGA cancer types, although the frequency peak of the SoMAS group shifted to different directions in different cancers (Figure S6A). While in most cancers, the HMP frequency peak of the SoMAS group shifted right, implying the SoMAS pairs exhibited more significant correlation in chromatin accessibility compared to non-SoMAS pairs, the HMP frequency peak of the SoMAS group in LUAD shifts dramatically to the left (Figure 6F, left). This means that in LUAD, the generally high co-occurrence of chromatin access in gene pairs was largely reduced in the somatic mutation associated alternative splicing events, which represents a special case relative to other cancer types. We further confirmed this kind of peak shift was absent if the compared groups were picked from the same pools (i.e., both SoMAS, or both non-SoMAS pairs) (Figure S6B-C; Figure 6F, middle and right). These results corroborate that the co-profile of chromatin accessibility around SoMAS genes and targets is largely disrupted, compared to general, randomly chosen gene pairs (Figure 6G). These ATAC-seq data together imply that somatic mutations appear to manifest biophysical differences in local chromatin accessibility at both the site of the mutation and the protein’s target sites. Therefore, it brings some new insights regarding the impact of somatic mutations on gene isoform expression potentially mediated by local chromatin accessibility.

### SoMAS genes are well supported by independent cohorts and methods

While currently the literature lacks a comprehensive mutation vs. isoform expression association study like SoMAS, there indeed exists piecemeal investigations of SNV-AS associations on particular types of SNVs or AS events that well support the SoMAS results. For example, an integrative analysis based on multi-omics data from six TCGA cancers identified 1,030 genes harboring somatic exonic SNVs (seSNVs) that potentially disrupt the gene-splicing process, including intron retention and exon skipping events [34]. SoMAS confirmed that 83 of these genes are associated with differential isoform expression in one or more of the six cancers (Table S7). Another study based on data from all 33 TCGA cancer types obtained 1,607 genes harboring splice-site-creating mutations (SCMs) [13], of which 103 genes were detected as SoMAS genes in various cancers (Table S7).

Strikingly, 10 out of the 14 (71.4%) confirmed TSGs with SNVs causing intron retention [34] were identified as SoMAS genes across different cancers; while 14 out of the 16 (87.5%) highlighted genes with recurrent SCMs [13] were detected as SoMAS genes, generally in multiple cancers (Figure 7A). A more general, independent analysis of splicing profiles using the same 33 TCGA cancers identified seven *trans*-sQTLs in six unique genes [14]. Four of them (66.7%), including *SF3B1*, *IDH1*, *EGFR* and *PPP2R1A*, are discovered by SoMAS in a varied number of cancer types (Figure 7A).

**Figure 7.**
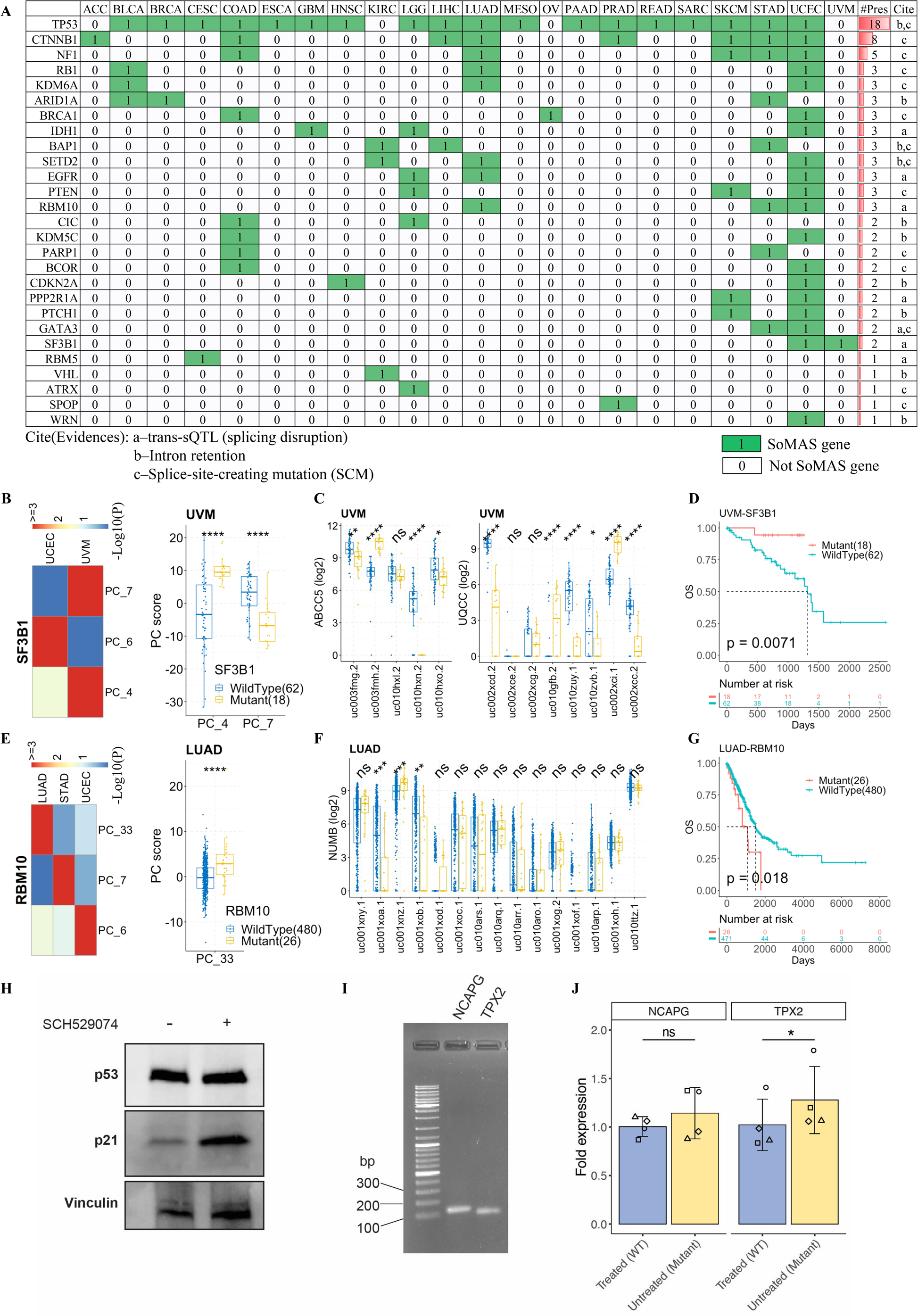
SoMAS genes are well evidenced by independent studies. (A) A catalogue of SoMAS genes that are explicitly supported by previous research. a: genes with *trans*-sQTLs that were reported to disrupt splicing by (Kahles et al., 2018); b: genes with SNVs causing intron retention by (Jung et al., 2015); c: genes with splice-site-creating mutations (SCMs) by (Jayasinghe et al., 2018). (B) SoMAS analysis results for SF3B1 in TCGA cancers. (C) Differential isoform expression analysis on UQCC and ABCC5 based on SF3B1 mutation status in UVM cancer. (D) Survival analysis based on the mutation status of SF3B1 in UVM cancer. (E) SoMAS analysis results for RBM10 in TCGA cancers. (F) Differential isoform expression analysis on NUMB based on RBM10 mutation status in LUAD cancer. (G) Survival analysis based on the mutation status of RBM10 in LUAD cancer. (H) Western blot generated from NCI-H1975 cell lysates. Cells were treated with 6µM SCH529074 compound or mock (DMSO) for 24 hours. Expression of p53 and p21 is shown. Vinculin was included as a loading control. (I) Gel electrophoresis of PCR products generated from NCI-H1975 cDNA. Primers targeting either NCAPG (isoform uc003gpp.2, lane 2) or TPX2 (isoform uc002wwp.1, lane 3) were tested. Lane 1 shows the DNA ladder. (J) Relative expression of the NCAPG isoform uc003gpp.2 (left) or the TPX2 isoform uc002wwp.1 isoform (right) in NCI-H1975 cells either treated with SCH529074 compound to restore WT function of p53 or left untreated, and thus carrying the R273H mutated p53. Expression level was measured by qRT-PCR. The height of the bars shows the mean expression level across 4 biological replicates (each assigned a symbol) and the error bars show the standard deviation from the mean. Expression was calculated for the untreated cells relative to the mean of the treated cells (normalized to 1). Significance levels were derived from one-sided paired t-test. Wilcoxon rank-sum test or t-test for differential analysis, \*\*\*\**P*<1e-4, \*\*\**P*<1e-3, \*\*\**P*<0.01, \**P*<0.05, ns: non-significant. See also Table S7 and Figure S7.

Among the genes with splicing-associated mutations, the splicing factor 3B subunit 1 (*SF3B1*) was consistently reported to be associated with differential alternative splicing of several protein-coding and noncoding genes in human cancers [14–16]. Particularly, AS events in *ABCC5* and *UQCC* have been validated as consequences of *SF3B1* mutations by qRT-PCR in UVM (uveal melanoma) cancer. Meanwhile, *SF3B1* mutations are relevant to patient survival, and are associated with better and worse prognosis in UVM [15] and CLL (chronic lymphocytic leukemia) [35] cancers, respectively. While the TCGA data does not cover the CLL cancer, our pipeline detected *SF3B1* as a SoMAS gene in UVM and UCEC cancers (Figure 7B), and accurately predicted ABCC5 and UQCC as its targets (ranking 150 and 88 in the unique gene list respectively) in UVM (Figure 7C). Another two SF3B1 targets predicted by SoMAS in UVM, ADAM12 and GAS8 (growth arrest-specific 8), despite marginal significance level (Figure S7A), have also been validated by the qRT-PCR assay. SoMAS further identified two novel GAS family genes, GAS7 and GAS6, as SF3B1 targets with high confidence (GAS7 is marginally significant), reported for the first time here. The targets of SF3B1 in UCEC are quite different than those in UVM, like other SoMAS genes, and ABCC5 and UQCC are not top (although ABCC5 tests marginally significant) targets of SF3B1 in UCEC (Figure S7B). In addition, we corroborated that patients with mutant SF3B1 have a better survival than those with wildtype SF3B1 in both UVM and UCEC cancers (Figure 7D, Figure S7C).

Additional published studies of relevant cancer genes support our SoMAS findings. For example, the RNA-binding motif (RBM) protein family genes, including RBM5, RBM6 and RBM10, have been shown to regulate splicing of NUMB to promote lung cancer cell growth [17]. SoMAS detected RBM5 as a SoMAS gene in CESC cancer, and RBM10 in three cancer types, including LUAD, STAD and UCEC (Figure 7E). SoMAS also predicted different isoforms of NUMB as the RBM10 targets in different cancers (Figure 7F, Figure S7D, F). Consistent with RBM10 function in promoting cell proliferation in lung cancer, LUAD patients with RBM10 mutations showed a significantly worse prognosis than those with wildtype RBM10 (Figure 7G). However, in STAD and UCEC cancers SNVs on RBM10 displayed a strong protective effect against cancer (Figure S7E, G), a phenomenon that gained increasing attention recently [36]. Taking these data together, we conclude that the biological and clinical relevance of the SoMAS genes are strongly supported by previous studies using independent methodologies and/or patient cohorts.

To validate the findings in an experimentally controlled setting, we sought to establish a cell-based assay that would allow comparison of isoform expression between a wild type (WT) and mutant cell line. With a focus on *TP53* and its role in lung cancer (Figure 2H), we tested the lung adenocarcinoma cell line NCI-H1975 carrying the *TP53* mutation R273H. This mutation occurs in the DNA binding domain affecting the ability of the protein to bind the p53-responsive element [37]. Treatment of mutant NCI-H1975 cells with compound SCH529074 is known to restore p53 DNA binding capacity, and caused an upregulation of downstream targets of p53, such as p21 [38], compared to mock treatment with DMSO (Figure 7H).

Having confirmed the action of the SCH529074 compound, we next measured the expression levels of isoforms associated with *TP53* mutation status. Two isoforms among the top 15 (ranked by PC loading in the SoMAS analysis) (Figure 2H and Table S7C) were tested by qRT-PCR at unique exon-exon junctions: *NCAPG* isoform uc003gpp.2 that contains a unique exon (exon 5) and *TPX2* isoform uc002wwp.1 that skips exon 10 and creates a unique exon junction between exons 10 and 12 (Figure S7H). Expected sizes of 127 bp for *NCAPG* and 99 bp for *TPX2* were obtained, indicating that only the specific isoforms of interest were amplified (Figure 7I).

We then quantified the isoform expression levels in the NCI-H1975 cells treated with 6µM SCH529074 compound or DMSO by qRT-PCR. Measurement entailed a delta-delta Ct analysis with beta-Actin acting as the housekeeping gene. *NCAPG* isoform uc003gpp.2 showed a slight, overall higher normalized expression when cells retained the non-functional p53 protein (mutant) compared to the treated controls which reverted p53 to wildtype (Figure 7J, left). However, no statistical significance was found. On the other hand, all four experiments of *TPX2* isoform uc002wwp.1 showed higher normalized expression in the *TP53* mutant group compared to the wildtype (average fold increase: 1.28, P-value = 0.02, one-sided, paired t-test) (Figure 7J, right). These wet-lab experimental data represent a proof-of-principle that further supports the SoMAS predictions.

## Methods

### TCGA data acquisition and preprocessing

Gene isoform expression data, gene somatic mutation profiles, and patient clinical information for 33 cancer types were downloaded from TCGA data portal (https://portal.gdc.cancer.gov/) with the R package TCGAbiolinks [39]. Specifically, the Illumina HiSeq RNASeqV2 RSEM-normalized isoform expression level for 73,599 gene isoforms were downloaded for each individual cancer type. These isoforms consist of 29,181 unique coding and noncoding human genes. When downloading the somatic mutation data, we chose the MUTECT2 pipeline which covers 22,029 different genes across all cancer types. We focused only on the single nucleotide variants (SNVs), which on average consists of 91.5% of all somatic mutations detected in each of the 33 TCGA cancer types (Table S1). For clinical information, the patient survival time, quantified as ‘days to death’ (overall survival time) and ‘days to last follow up’ (censored) were extracted and converted to numeric values for subsequent analysis. Sample type (tumor vs. adjacent normal) was determined by parsing the patient barcodes, and we focused on the tumor samples in most analyses unless otherwise noted. The specific tumor stage of each tumor sample was further extracted and categorized into four large groups I-IV. The tumor purity information was derived from a published consensus measurement of purity estimation (CPE) [40].

### ATAC-seq data acquisition and preprocessing

The matched ATAC-seq data for 23 out of 33 TCGA cancer types were downloaded from the NCI GDC page (https://gdc.cancer.gov/) in the form of cancer type-specific log2-transformed normalized ATAC-seq insertion count matrices (TCGA-ATAC_PanCan_Log2Norm_Counts.rds) [33]. A total of 796 individual technical replicates from 23 cancer types were obtained, with sample size ranging from 12 in CESC to 146 in BRCA (median = 29). The number of ATAC peaks ranged from 56,112 in CESC to 215,920 in BRCA with a median of 102,550. We further extracted the genomic coordinates and annotation type (including 3’ UTR, 5’ UTR, Distal, Exon, Intron and Promoter) of each ATAC peak for downstream analysis.

### RBP, SF and TF genes collection and filtering

We curated a total of 1,972 RNA-binding Protein (RBP) genes from MSigDB [26] and 404 splicing factor (SF) genes from published literature [16]. Among those, 1,681 RBP and 394 SF genes overlapping with the TCGA somatic mutation data were retained in the subsequent analysis. We downloaded transcription factors (TFs) and their targets with the R data package tftargets (https://github. com/slowkow/tftargets). This dataset includes human TF information curated from six published databases: TRED, ITFP, ENCODE, Neph2012, TRRUST, Marbach2016. As previously did [41], we mapped the Entrez gene IDs into gene symbols using two R packages: annotate and org.Hs.eg.db. After integration and removal of duplicates, we obtained 2,705 TFs with their target genes. We only focused on 2,478 TFs overlapping the TCGA somatic mutation data in the downstream analysis.

### Clustering of all TCGA samples based on gene isoform expression

To investigate the overall pattern of gene isoform expression across cancer types, we pooled the gene isoform expression data of all 33 TCGA cancer types together as a gene isoform-by-sample expression matrix for a clustering analysis. The original matrix includes 73,599 gene isoforms and 10,464 TCGA samples, covering 726 normal and 9,738 tumor samples. Genes expressed in less than 3 samples and samples expressing less than 200 isoforms were excluded for data quality control, after which we obtained a new expression matrix with 71,550 gene isoforms and 10,464 samples.

The R package Seurat [22] was applied to implement the clustering analysis and visualization. For the clustering based on gene isoform expression, we first normalized the expression level of each sample with a LogNormalize method as y = log(1+x/sum(x)*10,000), where x is the original 71,550-long vector of isoform expression for a sample in study, and log refers to the natural logarithm. Next, we identified 3,667 most variable gene isoforms (features) with mean.var.plot (mvp) method built in the Seurat package. The variable gene isoforms behave as outliers on a ‘mean variability plot’, and the mvp method detects those outliers while controlling for the strong relationship between variability and average expression. We centered and scaled the expression for each isoform by y = (x-mean(x))/sd(x), where x is the vector of expression along samples for an isoform and mean and sd refer to the average and standard deviation of expression for that isoform. Then we conducted a linear dimension reduction with principal component analysis (PCA) and reduced the dimension from 3,667 to 50. Finally, we implemented the clustering analysis based on the new 50-D meta-isoform expression matrix by calculating the 20-nearest neighbors and constructing the shared nearest neighbor (SNN) graph with Seurat built-in algorithms.

### Clustering of all TCGA samples based on gene somatic mutation

The workflow of clustering based on gene somatic mutation is similar to that based on the gene isoform expression, while the specific configurations are a little different. To generate the feature-by-sample matrix for gene somatic mutation, we counted the number of single nucleotide variants (or SNV for short, denoted as ‘SNP’ in the TCGA database) consisting of 13 different variant categories for each gene in each cancer type, including 3’-Flank, 3’-UTR, 5’-Flank, 5’-UTR, IGR, Intron, Missense_Mutation, Nonsense_Mutation, RNA, Silent, Splice_Region, Splice_Site, Translation_Start_Site. Each feature is marked as a ‘gene name + variant category’ combination. In this way, we annotated a total of 127,009 different mutation features for all 33 cancer types. After removing features present in less than 5 samples, we eventually obtained 76,731 mutation features, and accordingly a mutation matrix of 76,731*10,178. We further adjusted the initial counts of mutations by dividing the raw mutation counts in each gene in each cancer type by the length of that gene. Since mutation counts are more discrete compared to isoform expression, we normalized the mutation counts of each sample by the RC (relative counts) method as y = x/sum(x)*100, i.e., feature counts for each sample are divided by the total counts for that sample and multiplied by a scale factor (100). We chose the top 2,000 variable features with the local polynomial regression model (‘vst’ method built in Seurat) for the subsequent linear dimension reduction (PCA, also reduced to 50-D) and clustering analysis on the mutation data.

### Comparison and visualization of different clustering results

After the clustering analysis based on the gene isoform expression and gene somatic mutation profiles, we generated the cluster residence heatmap to compare the clustering results with the cancer tissue types of all the TCGA samples, as well as the published iCluster membership [21]. We further conducted a non-linear dimension reduction analysis based on the PCA output for both gene isoform expression and gene somatic mutation data, with Uniform Manifold Approximation and Projection for Dimension Reduction (UMAP) technique [42]. TCGA samples were mapped to the 2D UMAP space derived from gene isoform expression levels and colored in different ways, including Isoform and Mutation clusters, cancer tissue type (33 TCGA cancer types), sample type (tumor + normal), and clustering results based on other TCGA molecular data types from published work. The variation of information distance between two clusterings of the same objects was calculated with the vi.dist function in R package mmcluster.

### Principal component analysis (PCA) of gene isoform expression in individual cancers

The SoMAS pipeline was applied to the tumor samples of each cancer type. The preprocessing steps for the PCA in individual cancer types, including the selection of most variable isoforms with the ‘mvp’ method, normalization, and scaling of the original isoform expression data, are the same as that for the pooled samples as elucidated in above sections. As illustrated in Figure 2A, the only modification is that the ‘mvp’ method was performed on the 59,866 transcripts derived from 15,448 multi-isoform genes instead of the original 73,599 transcripts from 29,181 gene loci. Following the preprocessing steps, we transformed the informative gene isoform expression matrix into a low-dimensional PC score matrix with principal component analysis (PCA) before the subsequent differential analysis. Specifically, denote the preprocessed gene isoform expression matrix and the output PC score matrix as ***E*** and **S** respectively as follows,

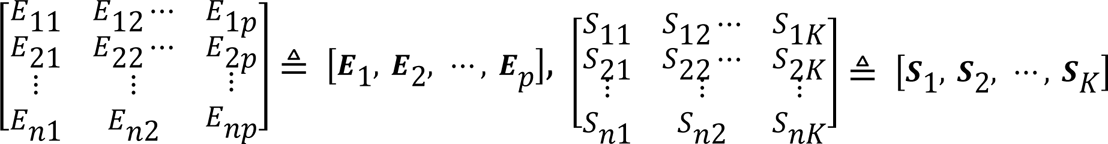

Where *n*, *p* and *K* stand for the number of samples, the number of gene isoforms (transcripts) and the number of PC-axes (or the dimension of the new space) respectively. The PCA algorithm tries to calculate the PC loading matrix

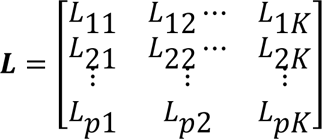

such that ***E*** × ***L*** = ***S***. In other words, ***S*** is linearly transformed from ***E***, and ***L*** represents the linear transformation that stores the corresponding combination coefficients. Taking the first PC-axis (PC_1) as an example, from the basic matrix multiplication operation it yields that ***S***_1_ = 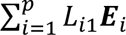, which means that the PC score for all *n* samples along PC_1 coordinate is a linear combination of the expression level of all original *p* gene isoforms. Therefore, ***S***_1_ is expected to be significantly correlated (either positively or negatively) with those ***E***_i_’s with largest coefficients *L*_i1_’s (either positive or negative). In this sense, the significant association between the mutation status of a gene and the PC_1 score is equivalent to the significant association between the gene mutation and those top isoforms (with largest *L*_i1_’s). It should be noted that this logic holds across all other PC coordinates, and this makes the mathematical foundation of that the SoMAS pipeline can identify all genes significantly associated with a mutant gene at one time by performing the differential PC score analysis instead of the direct differential gene isoform expression analysis. Throughout the work, we denote the top 100 significantly associated isoforms and their corresponding genes as the targets of the SoMAS gene (or the gene mutation in some contexts) unless otherwise stated.

### Differential analysis and visualization

We performed three types of differential analyses in this work, including differential PC score (from SoMAS), differential isoform expression (from RNA-seq), and differential chromatin accessibility (from ATAC-seq) analyses. In all analyses, samples were first divided into two groups based on the mutation status (wildtype vs. mutant) of a particular gene. Then a Wilcoxon rank-sum test was applied to compare the PC score or gene isoform expression between the two groups. For chromatin accessibility, *t*-test instead of Wilcoxon test was used due to smaller sample sizes. P-values of both Wilcoxon and *t*-test were provided as \**P*<0.05, \*\**P*<0.01, \*\*\**P*<1e-3, and \*\*\*\**P*<1e-4 unless otherwise stated. The Harmonic mean p-value (HMP) was calculated to measure the combined significance for genes with multiple ATAC-seq sites, with HMP<0.05 deemed significant. The ggboxplot function in the R package ggpubr was employed to visualize the differential values between groups.

### Detection of dual-direction associations between SoMAS gene mutation and SoMAS target expression

A dual-direction association refers to the scenario that the mutation status of a SoMAS gene is significantly positively associated with at least one isoform of its target gene, and at the same time, negatively associated with at least one isoform of the same target gene, both at the P<0.05 level (i.e., |Log2FC|>0 and Pval<0.05) with Wilcoxon rank-sum test. Only genes representing the top 100 target isoforms of each SoMAS gene in each cancer type were considered. For each SoMAS gene in each cancer type, we counted the number of its target genes (falling their top 100 target isoforms) that have a dual-direction association with this SoMAS gene and denoted it as N.Dual.sig.top100. For each SoMAS gene in each cancer type, there can be one or more dual-direction associated genes. For each dual-direction associated gene, we first calculated the HMP (Harmonic mean p-value) of the p-values related to all isoforms having a significant association (Wilcoxon rank-sum test P<0.05) with the mutation status of that SoMAS gene, then took an average over those HMP values to generate an mHMP value for that SoMAS gene. This mHMP value is finally log10-transformed [-log10(mHMP)] to Log10mHMP to indicate the overall significance level of dual-direction association for that SoMAS gene.

### Correlation between gene isoform expression and local chromatin accessibility

The ATAC-seq sites overlapping a specific gene isoform were first determined according to their genomic coordinates. Then the correlation between the chromatin accessibility (measured by ATAC-seq) on each site and the expression level of this gene isoform (measured by RNA-seq) was quantified by a Kendall correlation analysis. The correlation coefficients and significant levels (p-values) were provided. The Harmonic mean p-value (HMP) was calculated to measure the combined significance for genes with multiple ATAC-seq sites, with HMP<0.05 deemed significant.

### Calculation of inter-cancer overlap of SoMAS genes and SoMAS targets

The overlap of SoMAS genes between two cancer types was measured by the Szymkiewicz– Simpson coefficient [43], which is defined as the size of intersection divided by the smaller of the size of the two sets: overlap(X, Y) = |X∩Y|/min(|X|, |Y|). Briefly, we calculated the overlap as the number of SoMAS genes detected in both cancer types divided by the number of SoMAS genes detected in the cancer type with less SoMAS genes. The overlap of SoMAS targets between two cancer types was determined with three steps: first, the overlapping SoMAS genes between these two cancer types were picked; second, for each overlapping SoMAS gene, we focused on the top 100 target isoforms of this SoMAS gene in each cancer type and the number of overlapping targets was calculated; third, this number of overlapping targets were averaged over all the overlapping SoMAS genes between the two cancer types, which was eventually designated as the overlap of SoMAS targets between the involved two cancer types.

### Signaling pathway and hallmark gene set enrichment analysis

We downloaded the latest 343 KEGG signaling pathways (as of March 2021) with the R package clusterProfiler [44], and 50 hallmark gene sets via the Molecular Signature Database (MSigDB) [26]. For simplicity, the member genes of all pathways/gene sets were trimmed to the background 22,029 genes covered by the TCGA somatic mutation data. In both pan-cancer and cancer-specific scenarios, we performed enrichment analysis of the genes of interest against each of the 343 KEGG signaling pathways and 50 hallmark gene sets separately with hypergeometric test. Specifically, suppose we have *q* genes of interest, to determine their enrichment against a particular pathway (gene set) **S** with *m* member genes (out of *N* background genes), if *k* out of the *q* genes hit the pathway, we can calculate the probability of observing *k* or more genes hitting the pathway as

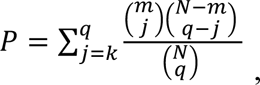

which gives the p-value of the hypergeometric test measuring the enrichment of the genes in this pathway or gene set under investigation.

### Survival analysis

To study the clinical relevance of a SoMAS gene and its corresponding PC score as single or combined variables, we performed both univariate and multivariate cox regression analysis on the two associated variables with the R package survivalAnalysis. The cox regression model quantifies the association between the two variables (individually or in combination) and patient survival time. We assessed the additive effect of the two variables in comparison with each individual by comparing the p-value for the same variable derived from univariate and multivariate cox regression analysis with paired t-test. We further confirmed the additive effect by directly comparing the overall survival rate between patient groups stratified by the combination of the two variables. Specifically, tumor samples were grouped in three different ways: (1) samples were divided into SoMAS gene wildtype and mutant groups; (2) samples were divided into positive and negative PC score groups; and (3) samples were divided into four groups based on the combination of the two variables (wildtype + positive, wildtype + negative, mutant + positive, mutant + negative). The survival difference was then quantified with log-rank test and visualized with Kaplan-Meier plot.

### Cell Culture and treatment with the SCH529074 compound

NCI-H1975 cells were obtained from ATCC and grown in DMEM (Gibco, Cat. No. 105660916), 10% Fetal Bovine Serum and 100 U/mL Penicillin-Streptomycin (Life Technologies, Cat. No. 15140122) at 37°C and 5% CO_2_. Cells were initially seeded at 0.5 x 10^5^ cells per well in a 6 well plate and grown overnight. Culture medium was replaced with medium containing 6 µM SCH529074 compound (MedChemExpress, Cat. No. HY-110088) or an equivalent volume of DMSO (mock treatment). Cells were grown for an additional 24 hours and harvested for protein or RNA.

### Protein extraction and western blot

Complete lysis buffer was prepared by combining 5 ml of NP-40 lysis buffer (50 mM Tris pH 7.0, 10 mM MgCl_2,_ 150mM NaCl, 1% NP-40), 1 cOmplete^™^ Mini EDTA-free Protease Inhibitor Cocktail tablet (Roche Cat. No. 04693159001), 50 µl phosphatase inhibitor cocktail 2 (Sigma, Cat. No. P5726), and 50 µl phosphatase inhibitor cocktail 3 (Sigma, Cat. No. P0044). Cells were pelleted, washed once in PBS, and lysed in 60-150 µL, depending on pellet size, then incubated on ice for 10-15 minutes. Lysates were centrifuged at 20,000 x g, 4°C for 10 min to remove cell debris. Supernatants were stored at −30°C. Protein concentration was determined by BCA assay (Pierce, Cat. No. 23225). For each gel equivalent amounts of protein were diluted to 30 *μ*L with additional lysis buffer. 10uL 4X Laemmli buffer (Bio-Rad, Cat. No. 1610747) and 2.4 *μ*L 2-mercaptoethanol were added to each sample, which were then heated to 95°C for 5 minutes and cooled to room temperature before loading. Gel electrophoresis was performed using 4–15% Mini-PROTEAN® TGX™ gels (BioRad, Cat. No. 4561084) and Tris/Glycine/SDS Running Buffer (BioRad, Cat. No. <u>1610732</u>). Proteins were transferred to PDVF membranes using Trans-Blot Turbo Mini 0.2 µm PVDF Transfer Packs (BioRad, Cat. No. 1704156) and the BioRad Transblot system. Blots were blocked in 5% skimmed milk dissolved in a 1X TBS buffer (BioRad, Cat. No. 1706435) for 1 hour at room temperature, then probed overnight at 4°C with primary antibodies diluted in a 5% BSA/ 1X TBS-0.1% Tween 20 (TBS-T) solution using either Vinculin (E1E9V) (Cell Signaling, Cat. No. 13901) diluted 1:2000, p53 (7F5) (Cell Signaling, Cat. No. 2527) diluted 1:4000, or p21 Waf1/Cip1 (12D1) (Cell Signaling, Cat. No. 2947) diluted 1:4000. Blots were washed 3 times for 10 minutes at room temperature with 0.1% TBS-T and probed with secondary antibody, Anti-rabbit IgG HRP-linked Antibody (Cell Signaling, Cat. No. 7074) diluted 1:1000 in 0.1% TBS-T for 1 hour at room temperature. Blots were washed 3 times with TBS-T for 10 minutes at room temperature and then developed with Pierce SuperSignal™ West Femto Maximum Sensitivity Substrate (Thermo Fisher, Cat. No. 340940). Blots were visualized with a BioRad GelDoc system.

### RNA extraction and conversion to cDNA

RNA was prepared from cells that had been pelleted and washed once with PBS, using the Ambion PureLink RNA kit (Invitrogen, Cat. No. 12183020) according to kit directions, including an on-column DNAse digestion step. cDNA was generated using a Qiagen Quantitect kit (Cat. No. 205311) according to kit instructions.

### Isoform specific PCR primers

PCR primers were designed such that forward primers were spanning a unique exon-exon junction for the target the specific isoform of interest. For *NCAPG* (NM_022346, isoform uc003gpp.2) the forward primer spans the junction between exon 4 and 5 (5’-AAGCTGGCTTATCAGGTTT-3’), while the reverse primer resides within exon 6 (5’-GCTTCTGCATAGCTTGTTTC-3’). For *TPX2* (NM_012112, isoform uc002wwp.1) the forward primer spans the junction between exon 10 and 11 (5’-GGAGCAAGAAGGATGATATTAACCTGT-3’), while the reverse primer resides within exon 11 (5’-CACGGTGTTTGGTTTGCAGT-3’). All primers were ordered from Integrated DNA Technologies with standard desalting and resuspended in DNase free water to a concentration of 100µM and stored at −20°C.

### qRT-PCR assay validation

Validation of each assay was performed on cDNA generated from mock-treated H1975-NCI cells. To confirm that each assay amplified a product of the expected size, a regular PCR was performed using 200nM of each forward and reverse primer, 2X Platinum^TM^ SuperFi^TM^ II PCR Master Mix (Invitrogen, Cat. No. 12368010), 0.5µl cDNA template and DNase free water to an adjusted volume of 25µl. The PCR reaction was performed in the SimpliAmp^TM^ Thermal Cycler (Applied Biosystems) using the following cycling conditions: 98°C for 2 min., then 40 cycles of 98°C for 30 seconds, 60°C for 30 seconds, and 72°C for 30 seconds, and a final 72°C for 5 min. PCR products were diluted 1:10 before loading on a 1% agarose gel alongside GeneRuler DNA Ladder Mix (Thermo Scientific, Cat. No. SM0331).

### qRT-PCR assay for assessment of relative expression

qRT-PCR analysis was performed for *NCAPG* and *TPX2*. Primers targeting beta-Actin (ACTB) (NM_001101) were obtained from Origene (Cat. No. HP204660) and included as the reference gene. qRT-PCR was performed as singleplex reaction with three technical replicates for each condition. Each reaction contained 500nM of forward and 500nM of reverse primer, PowerUp SYBR Green Master Mix (2X) (ThermoFisher Scientific, Cat. No. A25741), 0.5µl cDNA template and DNase free water to an adjusted volume of 25µl. The PCR reaction was performed in the C1000 Touch^TM^ Thermal Cycler with a CFX96^TM^ Optical Reaction Module (BioRad) using the following cycling conditions: UDG activation at 50°C for 2 min., 95°C for 2 min., 40 cycles of 95°C for 15 seconds 60°C for 1 minute with signal acquisition at the end of the annealing step. To monitor uniformity of the amplified products, a melting profile was added. Relative expression was calculated using the Delta-Delta Ct method (2^−ΔΔCt^ method). In brief, Ct values for technical replicates were averaged, and delta Ct was calculated for each sample using the beta Actin as reference gene. Finally, delta-delta Ct was calculated using the untreated (mock) NCI-H1975 as reference samples. Thus, the delta-Ct values for the biological replicates were averaged to create a ‘control average’. Result from each of the samples (treated or untreated) are presented as a fold expression relative to this control average.

## Discussion

Somatic single nucleotide variants (SNVs) have been increasingly recognized to drive human diseases especially cancer by inducing alternative splicing (AS) and altering gene expression patterns at the transcriptome level [45, 46]. Many efforts have been devoted to identifying SNVs that disrupt splicing (called splicing mutations) and exploring their implications in human oncogenesis. However, the previous research focused on the direct regulation of splicing by SNVs occurring in flanking exons (for *cis*-association) or a small number of known splicing factors/regulators (for *trans*-association) but ignored SNVs located in many other RNA-binding proteins (RBPs) or splicing factors (SFs), and even more upstream regulators (e.g., TFs) of splicing machinery that function indirectly. In addition, this direct gene-by-gene association study involves a tremendous computational burden and easily brings false positives to exhaust all SNV-AS combinations. To address these challenges, we developed a computational pipeline, called SoMAS (Somatic Mutation associated with Alternative Splicing), that integrates matched DNA-seq and RNA-seq data to systematically investigate the effect of somatic SNVs on AS profiles across the whole genome in human cancers.

We have shown that SoMAS is more efficient and versatile than existing methods in identification of AS-associated SNVs. Since SoMAS implements differential analysis along a small number of PC coordinates (always ≤50) rather than along thousands of original transcript axes directly, it dramatically outperforms the existing models in efficiency. Furthermore, SoMAS searches for the SNV-impacted transcripts over the whole genome simultaneously, which means that it determines both the *cis*- and *trans*-regulatory SNVs at the same time, indicating its versatility compared to previous studies. Most importantly, SoMAS overcomes the limitation of the low mutation frequencies of most cancer genes by examining the association between an SNV and the overall genome-wide landscape of gene splicing, well reducing the false positive rate incurred by the traditional gene-by-gene association study.

Applying SoMAS to 33 TCGA cancer types individually, we detected 7,140 SoMAS genes in total, including 1,650 and 908 and SoMAS genes that are simultaneously present in at least two and three cancer types, respectively. The SoMAS genes included many widely known oncogenes and tumor suppressors, exemplified by the significant enrichment of SoMAS genes in the Cancer Gene Census (CGC) genes. This verifies the hypothesis that mutations on cancer-associated genes could function via disrupting splicing and transcription processes. The SoMAS genes also covered a considerable number of RBPs and SFs, especially those RNA-binding SFs, indicating that SoMAS can identify direct regulators of the splicing process as well. Additionally, many TFs were identified as SoMAS genes in various cancers, meaning that SoMAS genes can impact gene isoform expression as a transcription regulator. Both the top 908 SoMAS genes and their top target genes are significantly enriched in tumor growth and metastasis related processes. Some SoMAS genes better stratify patients in terms of survival rate when in combination with expression of their target transcripts (aggregated as a PC score). These results clearly demonstrate the biological and clinical significance of the SoMAS genes.

Most previous studies looked for the direct association between a somatic (or germline) SNV and an explicit AS event with real split-reads support, which requires a thorough reanalysis of the raw RNA-seq data from scratch to count the split-reads, followed by an elaborate quantification of the AS event, e.g., with a percent spliced in (psi) value [14]. SoMAS examines the association between an SNV and the expression profiles of all transcripts derived from multi-isoform genes that have been already well annotated and quantified in TCGA. SoMAS accurately identified many genes that have been proven to directly impact gene splicing by independent methods or data sources, including splicing factors (e.g. *SF3B1*, *RBM10*) that disrupt splicing of specific genes in particular cancers (*trans*-effect) [15–17], and genes with mutations that induce splice site creation nearby such as *TP53*, *GATA3*, *KDM6A*, *PTEN*, *SETD2*, *RB1* and *CTNNB1* in different cancers (*cis*-effect) [13]. Together, we documented that mRNA abundance at the transcriptome level retains essential information regarding the gene-splicing profiles, which enables SoMAS to identify those splicing mutations through associations with isoform-level expression without the splicing details. This is well illustrated by the many SoMAS genes whose mutation status is significantly associated with isoform expression in a dual-direction manner (Figure S2F-H, Table S2B). Therefore, while SoMAS is not designed to detect the association between SNV and AS events directly, the association between an SNV and isoform-level expression of a gene can bring many insights about the effects of the SNV on splicing of those associated genes. In this sense, SNVs on SoMAS genes might be better interpreted as the isoform-level expression quantitative trait loci, or isoQTLs [47].

To better interpret this point, it’s worth noting that SoMAS focuses only on the 59,866 (instead of the original 73,599) isoforms that were derived from multiple-isoform genes (Figure 2A). That is, the single-isoform genes were excluded from the very beginning. This is because single-isoform genes can be easily spotted by the traditional differential expression analysis at the gene level. Our study added value by screening differences across all isoforms, including both major and minor ones, to detect expression alterations. On the other hand, since each particular isoform comes from the full gene annotation, every isoform, no matter major or minor, comprises a subset of all the exons, which refines and specializes its activity versus the entire set of exons. In this sense, expression change of a particular isoform indicates some specialized functions of a gene are being altered. Hence, SoMAS can evaluate the impact of gene mutation on those specialized functions of its target genes.

The target isoforms of most SoMAS genes are distributed across the whole genome, generally proportional to the number of transcripts located in each chromosome. This implies that the *trans*-acting associations dominate the *cis*-acting associations. A *trans*-acting association suggests that the SoMAS gene might impact the specific isoform’s expression as a TF, SF, or other co-regulator of gene splicing and/or transcription. It’s also possible that the target in each *trans*-association acts as an intermediate expression regulator of downstream genes. Indeed, we have shown that many RBP and SF genes ranked the top targets of some SoMAS genes in multiple cancer types. The splicing factor *HNRNPK* gene represents such an example. Previous study showed that *TP53* mutations modify RNA splicing by promoting *HNRNPK* expression in pancreatic cancer [18]. SoMAS predicts *HNRNPK* as a top target of some SoMAS genes in 11 cancer types (Table S5). However, *TP53* was not present in any list of those SoMAS genes. This might be because our current pipeline includes all SNVs in the mutant group instead of only R172H/R175H as did in the literature. When we customized our pipeline to the mentioned genotype in PAAD cancer, we observed elevated HNRNPK expression in the TP53-mutant group compared to normal control with marginal significance level, which shows certain consistency with the literature. Hence, our pipeline is easy to be calibrated to determine association between designated genotypes and gene isoform expression patterns.

With matched ATAC-seq data we disclosed that both gene somatic mutation and gene isoform expression are correlated to local chromatin accessibility to some extent, especially in the intron and promoter regions (Figure 6A-C). This is comprehensible considering that the interplay between mutational process and 3-D genome organization has been reported [48]. The somatic mutation is expected to impact the binding of chromatin-binding factors to the chromosome, which is another possible mechanism of how the somatic mutation influences the nearby chromatin conformation and hence accessibility. The effect of chromatin accessibility on gene isoform expression is more straightforward and natural, in that the splicing and transcription processes need to recruit a series of factors/regulators on-site. We further revealed that the co-profile of chromatin accessibility around the SoMAS genes and predicted targets is largely disrupted compared to randomly chosen pairs. However, the mechanisms and implications underlying this observation remain to be elucidated.

Some potential confounding factors need to be clarified. First, the number of SoMAS genes detected in each cancer type has little correlation with the sample size (Figure 2B, upper). Second, although the correlation between the numbers of SoMAS genes and total qualifying mutant genes (mutational burden) across cancer types are statistically significant, that doesn’t confound the detection of SoMAS genes for two reasons: 1) the overall too few SoMAS genes detected in most cancers, i.e., no more than 10 SoMAS genes were detected in 18 out of 33 TCGA cancer types, making the impact of mutational burden negligible; 2) for each mutant gene to test, we performed a multiple test along the 50 PC coordinates and have corrected the p-value with Bonferroni method already (setting the threshold of minimum p-value as 0.05/50=0.001). Indeed, when we carried out a multiple test correction on all original p-values and set the threshold of adjusted p-value=0.05 as commonly used, we obtained much more SoMAS genes in most cancer types, which apparently were false positives. Third, the gene length does not increase the likelihood for a gene to be detected as a SoMAS gene despite a potentially higher mutation frequency among tumor samples. This is because SoMAS compares gene isoform expression between groups with and without mutation of a particular gene. In this sense, higher mutation frequency of this gene only impacts grouping of the samples but is independent of the expression level of each group. Due to this reason, we set a relatively low threshold of mutation frequency (2% of all tumor samples in each cancer type, but subject to a minimum of 5 samples) for each mutant gene to qualify for a test.

To conclude, SoMAS is a novel pipeline that can efficiently identify SNVs impacting gene splicing and transcription at the transcriptome level, and the pan-cancer SoMAS genes we identified proved to bear important biological and clinical meaning.

## Data availability

Data generated in this study are published with this work as supplementary materials. We also created a GitHub repository (https://github.com/elnitskilab/SoMAS) for this project, which includes all R codes, tutorials and demo datasets that demonstrate usage of the SoMAS pipeline and interpretation of its output.

## Acknowledgements

We thank all members of the Elnitski Lab for valuable discussions. This work was supported by the Intramural program of the National Human Genome Research Institute, National Institutes of Health (L.E., 1ZIAHG200323).

## Reference

1. Chen, M. and J.L. Manley, Mechanisms of alternative splicing regulation: insights from molecular and genomics approaches. Nat Rev Mol Cell Biol, 2009. 10(11): p. 741–54.

2. Hanahan, D. and R.A. Weinberg, Hallmarks of cancer: the next generation. Cell, 2011. 144(5): p. 646–74.

3. Oltean, S. and D.O. Bates, Hallmarks of alternative splicing in cancer. Oncogene, 2014. 33(46): p. 5311–8.

4. Holland, D.O., et al., Characterization and clustering of kinase isoform expression in metastatic melanoma. PLoS Comput Biol, 2022. 18(5): p. e1010065.

5. Vogelstein, B., et al., Cancer genome landscapes. Science, 2013. 339(6127): p. 1546–58.

6. Tan, H., J. Bao, and X. Zhou, A novel missense-mutation-related feature extraction scheme for ‘driver’ mutation identification. Bioinformatics, 2012. 28(22): p. 2948–55.

7. Tan, H., J. Bao, and X. Zhou, Genome-wide mutational spectra analysis reveals significant cancer-specific heterogeneity. Sci Rep, 2015. 5: p. 12566.

8. Tan, H., Somatic mutation in noncoding regions: The sound of silence. EBioMedicine, 2020. 61: p. 103084.

9. Cartegni, L., S.L. Chew, and A.R. Krainer, Listening to silence and understanding nonsense: exonic mutations that affect splicing. Nat Rev Genet, 2002. 3(4): p. 285–98.

10. Lopez-Bigas, N., et al., Are splicing mutations the most frequent cause of hereditary disease? FEBS Lett, 2005. 579(9): p. 1900–3.

11. Woolfe, A., J.C. Mullikin, and L. Elnitski, Genomic features defining exonic variants that modulate splicing. Genome Biol, 2010. 11(2): p. R20.

12. Jung, H., K.S. Lee, and J.K. Choi, Comprehensive characterisation of intronic mis-splicing mutations in human cancers. Oncogene, 2021. 40(7): p. 1347–1361.

13. Jayasinghe, R.G., et al., Systematic Analysis of Splice-Site-Creating Mutations in Cancer. Cell Rep, 2018. 23(1): p. 270–281 e3.

14. Kahles, A., et al., Comprehensive Analysis of Alternative Splicing Across Tumors from 8,705 Patients. Cancer Cell, 2018. 34(2): p. 211–224 e6.

15. Furney, S.J., et al., SF3B1 mutations are associated with alternative splicing in uveal melanoma. Cancer Discov, 2013. 3(10): p. 1122–1129.

16. Seiler, M., et al., Somatic Mutational Landscape of Splicing Factor Genes and Their Functional Consequences across 33 Cancer Types. Cell Rep, 2018. 23(1): p. 282–296 e4.

17. Bechara, E.G., et al., RBM5, 6, and 10 differentially regulate NUMB alternative splicing to control cancer cell proliferation. Mol Cell, 2013. 52(5): p. 720–33.

18. Escobar-Hoyos, L.F., et al., Altered RNA Splicing by Mutant p53 Activates Oncogenic RAS Signaling in Pancreatic Cancer. Cancer Cell, 2020. 38(2): p. 198–211 e8.

19. Bailey, M.H., et al., Comprehensive Characterization of Cancer Driver Genes and Mutations. Cell, 2018. 174(4): p. 1034–1035.

20. Lawrence, M.S., et al., Mutational heterogeneity in cancer and the search for new cancer-associated genes. Nature, 2013. 499(7457): p. 214–8.

21. Hoadley, K.A., et al., Cell-of-Origin Patterns Dominate the Molecular Classification of 10,000 Tumors from 33 Types of Cancer. Cell, 2018. 173(2): p. 291–304 e6.

22. Butler, A., et al., Integrating single-cell transcriptomic data across different conditions, technologies, and species. Nature Biotechnology, 2018. 36(5): p. 411-+.

23. Ringner, M., What is principal component analysis? Nat Biotechnol, 2008. 26(3): p. 303–4.

24. Jolliffe, I.T. and J. Cadima, Principal component analysis: a review and recent developments. Philos Trans A Math Phys Eng Sci, 2016. 374(2065): p. 20150202.

25. Sondka, Z., et al., The COSMIC Cancer Gene Census: describing genetic dysfunction across all human cancers. Nat Rev Cancer, 2018. 18(11): p. 696–705.

26. Liberzon, A., et al., The Molecular Signatures Database (MSigDB) hallmark gene set collection. Cell Syst, 2015. 1(6): p. 417–425.

27. Licatalosi, D.D. and R.B. Darnell, RNA processing and its regulation: global insights into biological networks. Nat Rev Genet, 2010. 11(1): p. 75–87.

28. Hug, N. and J.F. Caceres, The RNA helicase DHX34 activates NMD by promoting a transition from the surveillance to the decay-inducing complex. Cell Rep, 2014. 8(6): p. 1845–1856.

29. Wu, D., et al., MZF1 mediates oncogene-induced senescence by promoting the transcription of p16(INK4A). Oncogene, 2022. 41(3): p. 414–426.

30. Rao, V.N., et al., Analysis of the DNA-binding and transcriptional activation functions of human Fli-1 protein. Oncogene, 1993. 8(8): p. 2167–73.

31. Vandin, F., E. Upfal, and B.J. Raphael, De novo discovery of mutated driver pathways in cancer. Genome Res, 2012. 22(2): p. 375–85.

32. Group, P.T.C., et al., Genomic basis for RNA alterations in cancer. Nature, 2020. 578(7793): p. 129–136.

33. Corces, M.R., et al., The chromatin accessibility landscape of primary human cancers. Science, 2018. 362(6413).

34. Jung, H., et al., Intron retention is a widespread mechanism of tumor-suppressor inactivation. Nat Genet, 2015. 47(11): p. 1242–8.

35. Cazzola, M., et al., Biologic and clinical significance of somatic mutations of SF3B1 in myeloid and lymphoid neoplasms. Blood, 2013. 121(2): p. 260–9.

36. Tan, H., On the Protective Effects of Gene SNPs Against Human Cancer. EBioMedicine, 2018. 33: p. 4–5.

37. Parrales, A. and T. Iwakuma, Targeting Oncogenic Mutant p53 for Cancer Therapy. Front Oncol, 2015. 5: p. 288.

38. Demma, M., et al., SCH529074, a small molecule activator of mutant p53, which binds p53 DNA binding domain (DBD), restores growth-suppressive function to mutant p53 and interrupts HDM2-mediated ubiquitination of wild type p53. J Biol Chem, 2010. 285(14): p. 10198–212.

39. Colaprico, A., et al., TCGAbiolinks: an R/Bioconductor package for integrative analysis of TCGA data. Nucleic Acids Research, 2016. 44(8).

40. Aran, D., M. Sirota, and A.J. Butte, Systematic pan-cancer analysis of tumour purity. Nat Commun, 2015. 6: p. 8971.

41. Tan, H., et al., Pan-cancer analysis on microRNA-associated gene activation. EBioMedicine, 2019.

42. McInnes, L., J. Healy, and J. Melville, UMAP: Uniform Manifold Approximation and Projection for Dimension Reduction, in arXiv preprint arXiv. 2018.

43. Vijaymeena, M.K. and K. Kavitha, A Survey on Similarity Measures in Text Mining. Machine Learning and Applications, 2016. 3(1): p. 19–28.

44. Yu, G., et al., clusterProfiler: an R package for comparing biological themes among gene clusters. OMICS, 2012. 16(5): p. 284–7.

45. Baralle, D. and M. Baralle, Splicing in action: assessing disease causing sequence changes. J Med Genet, 2005. 42(10): p. 737–48.

46. Garcia-Blanco, M.A., A.P. Baraniak, and E.L. Lasda, Alternative splicing in disease and therapy. Nat Biotechnol, 2004. 22(5): p. 535–46.

47. Gandal, M.J., et al., Transcriptome-wide isoform-level dysregulation in ASD, schizophrenia, and bipolar disorder. Science, 2018. 362(6420).

48. Akdemir, K.C., et al., Somatic mutation distributions in cancer genomes vary with three-dimensional chromatin structure. Nat Genet, 2020. 52(11): p. 1178–1188.

